# High-efficiency Kemp eliminases by complete computational design

**DOI:** 10.1101/2025.01.04.631280

**Authors:** Dina Listov, Eva Vos, Gyula Hoffka, Shlomo Yakir Hoch, Andrej Berg, Shelly Hamer-Rogotner, Orly Dym, Shina Caroline Lynn Kamerlin, Sarel J. Fleishman

## Abstract

We present a fully computational workflow for *de novo* design of efficient enzymes using backbone fragments from natural proteins and without recourse to iterative experimental optimization. The best designed Kemp eliminase exhibits >140 mutations from any natural protein, high stability (>85 °C) and unprecedented catalytic efficiency (12,700 M^-1^s^-1^), surpassing previous computational designs by two orders of magnitude. We find that mutations both inside and outside the active site contribute synergistically to the high observed activity and stability. Mutation of an aromatic residue used in all prior Kemp eliminase designs increases efficiency to >10^5^ M^-1^s^-1^. Our approach addresses critical limitations in design methodology, generating stable, high-efficiency, new-to-nature enzymes in complex folds and enables testing hypotheses on the fundamentals of biocatalysis through a limited experimental effort.

## Main

Natural enzymes are exceptionally versatile, selective, and highly efficient catalysts. Yet, computational design of enzymes that match this proficiency, particularly for nonnatural reactions, remains elusive. Recent advances in computational design have enabled rapid and effective optimization of natural enzyme stability, expressibility, catalytic rate, and selectivity through fully computational workflows (*1–4*). Furthermore, advances in fold design enabled grafting natural or engineered active sites into idealized *de novo* backbones (*5, 6*). By contrast, enzymes designed *de novo*, that is, without recourse to naturally occurring enzymes that catalyze the same reaction, are orders of magnitude less active relative to comparable natural ones (*7–11*). Previous studies have therefore employed repeated cycles of laboratory evolution, involving high-throughput screening of mutants, to reach effective enzymes (*11–15*). Such cycles are inefficient and restrict the application of *de novo* design to reactions that can be assayed in medium-to-high throughput. Critically, continuing to rely on screening large libraries and random mutations suggests that our understanding and control of the fundamentals of biocatalysis are far from complete.

The Kemp elimination (KE) reaction (Fig. 1A), a prototype for natural base-catalyzed proton abstraction, has long served as a model for studying *de novo* enzyme design, as no naturally occurring enzyme is known to catalyze this reaction. Despite increasing sophistication in protein design methods, computationally designed KEs exhibited low efficiencies (*k*_cat_*/K*_m_ 1-420 M^-1^s^-1^, *k*_cat_ 0.006-0.7 s^-1^) (*7, 9*) that are well below the efficiencies of the median natural enzyme (*k*_cat_/*K*_m_ 125,000 M^-1^s^-1^, *k*_cat_ 10 s^-1^) (*16*). These were subsequently optimized by laboratory evolution to 250,000 M^-1^s^-1^ (*12*). Analysis of these optimized enzymes confirmed that efficient catalysis depended on high shape complementarity between the ligand and active site and on precise positioning of the catalytic constellation (*12*) in a desolvated active-site environment (*13*). Nonetheless, these insights alone have not led to complete computational design of high-efficiency enzymes.

**Fig. 1.**
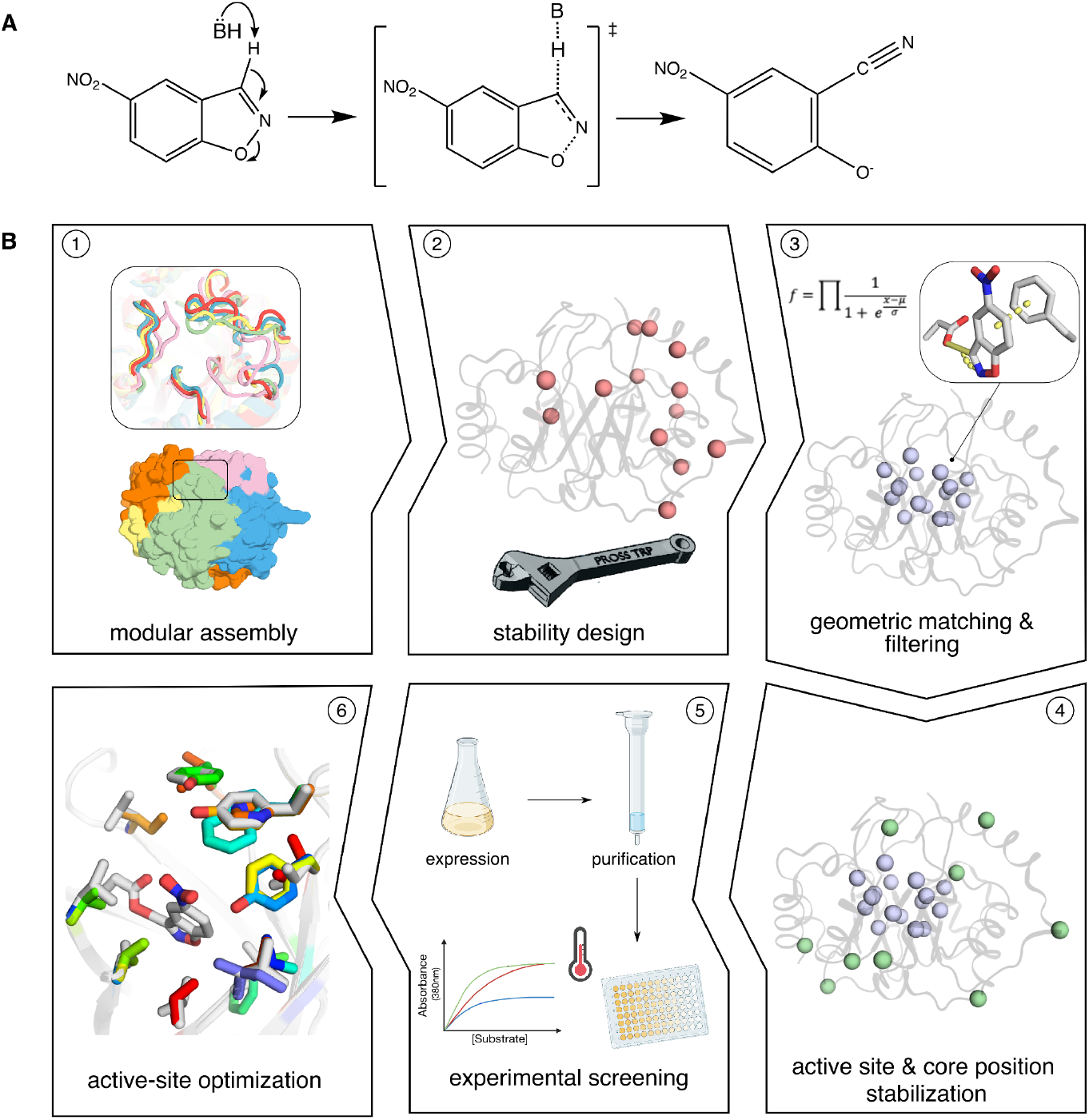
Key steps in the design workflow. **A**. Kemp elimination of 5-nitrobenzisoxazole. “B” is a base, implemented as the sidechain of Asp or Glu. **B**. Thousands of backbones are generated through combinatorial backbone assembly (step 1) and stabilized using PROSS (*3*) (step 2). Geometric matching (*37*) and active-site optimization with Rosetta yield millions of designs that are filtered by balancing energy terms that contribute to stability and activity (step 3). A few dozen top designs are chosen for further core and active-site stabilization using FuncLib (*4*) and pSUFER (*40*) (step 4). Following experimental screening (step 5), we apply FuncLib to the active sites of select functional designs (step 6).

The limitations of *de novo* enzyme design methodology have been the subject of intense study (*17–20*). These analyses revealed that the designed active sites exhibited significant structural distortions relative to the design conception (*17, 20*). Notably, catalysis is extremely sensitive to molecular details, and shifts of the catalytic constellation by a few degrees or tenths of an Ångstrom from optimality may translate into orders of magnitude decreases in efficiency (*21*). Furthermore, designs often exhibited low stability and expressibility (*13*), limiting their ability to accommodate activity-enhancing mutations (*2, 13*). Additional concerns were that fixed-backbone design methods fail to precisely position nonnative catalytic groups (*7*); the molecular details of the designed transition state (theozyme) were uncertain (*12, 22*); and that protein dynamics (*23*) and long-range electrostatic interactions may be necessary to achieve high catalytic efficiency but are unaccounted for in the design process (*12, 24, 25*). Given these manifold concerns, *de novo* design of efficient enzymes clearly provides a fertile testing ground for hypotheses on biocatalysis and its relationship to protein folding, stability and activity. Here, we sought to develop a systematic strategy for *de novo* enzyme design without high-throughput screening and mutational randomization to probe the necessary and sufficient conditions for effective enzyme design.

### Encoding stability, foldability, and activity in enzyme design

Our working hypothesis is that effective enzyme design requires control over all protein degrees of freedom to establish stability, foldability, and accurate positioning of the theozyme. Foldability, the ability of the protein to fold uniquely into the design conception, has been a long-standing challenge for *de novo* enzyme design. Over the past decade, foldability has been partly addressed through *de novo* fold design, enabling the generation of numerous stable and accurately designed proteins (*26–29*). Such designs, however, overwhelmingly comprise simple topologies (*1*) and are unlikely to be versatile enough to accurately encode the catalytic constellations required for sophisticated chemical transformations. To date, no *de novo* fold has shown catalytic rates comparable to those of natural enzymes. Given these limitations, we decided to focus our efforts on the TIM-barrel fold, which is one of the most prevalent enzyme folds (*30, 31*). In this fold, the residues of the central β barrel are oriented towards the active-site cavity, providing many opportunities to optimize the placement of catalytic and substrate-binding groups. Therefore, notwithstanding the challenges seen in designing accurate and functional TIM barrels (*32, 33*), this fold stands out as an attractive framework for engineering new enzymatic functions.

We developed a computational method that can be applied, in principle, to any reaction, given a precomputed theozyme. The workflow starts by generating thousands of backbones using combinatorial assembly and design (Fig. 1B step, 1), which combines fragments from homologous proteins to generate new backbones (*34–36*) and applies PROSS design calculations to stabilize the designed native state (*3*) (Fig. 1B step, 2). The resulting structures display structural differences within the active-site pocket that may increase the likelihood of obtaining foldable backbones that position the theozyme and supporting residues in a catalytically competent and energetically relaxed constellation. Following backbone generation, we implement geometric matching (*37*) to position the KE theozyme (*7*) in each of the designed structures and optimize the remainder of the active site using Rosetta atomistic calculations (*38*) (Fig. 1B, step 3). These steps result in millions of designs that are filtered using an optimization objective function that balances potentially conflicting goals (*39*) that are critical for design of function, such as low system energy and high desolvation of the catalytic base. Selecting a few dozen top-scoring designs, we next stabilize the active site and positions in the protein core using FuncLib (*4*) and pSUFER (*40*) (Fig. 1B step, 4), resulting in designs with >100 mutations from any natural protein. Unlike previous approaches, this workflow emphasizes stability across the entire protein and capitalizes on the ability to generate thousands of stable, natural-like TIM-barrels that exhibit structural diversity in the active site (*35*).

### High-efficiency, stable, and accurate Kemp eliminases

We applied our pipeline to the indole-3-glycerol-phosphate synthase (IGPS) enzyme family, which can sterically accommodate the 5-nitrobenzisoxazole substrate and was previously used to design KEs (*7, 13*). The theozyme builds on a catalytic constellation derived from quantum-mechanical computations (*41, 42*). It includes a nucleophile, such as Asp or Glu, that serves as a base for proton abstraction from the substrate (5-nitrobenzisoxazole), and an aromatic sidechain that forms π-stacking interactions with the substrate in the transition state (Fig. 1B step 3). The latter interaction has been used in all computational KE studies to date for its ability to promote binding to the aromatic benzisoxazole rings (*7, 9*). Typical design studies also introduced a polar interaction with the isoxazole oxygen to stabilize the developing negative charge in the transition state (*7, 9*). We excluded this interaction from our theozyme because a water molecule can satisfy this requirement, and a misplaced polar group could reduce reactivity by lowering the p*K*_a_ of the catalytic base.

We selected 73 designs for experimental testing. The designs ranged from 245 to 268 amino acids and were diverse, with 30-93% sequence identity to one another and 41-59% identity to any natural protein. 66 designs expressed solubly and 14 showed cooperative thermal denaturation (fig. S1). Three designs showed measurable KE activity in an initial screen, with the top two designs, Des27 exhibiting an apparent *T*_m_ 79°C, *K*_m_ 0.5 mM, *k*_cat_ 0.07 s^-1^ and *k*_cat_*/K*_m_ 130 M^-1^s^-1^ and Des61 exhibiting *T*_m_ 65°C, *K*_m_ 1.3 mM, *k*_cat_ 0.3 s^-1^ and *k*_cat_*/K*_m_ 210 M^-1^s^-1^ (fig. S2, table S1, S2).

The catalytic rate and efficiency of the these designs are on a par with previously designed enzymes (*7, 9*), falling short by several orders of magnitude from comparable natural eliminases and from KEs that were subjected to laboratory evolution campaigns (*12, 13*). To optimize these designs computationally, we applied FuncLib to active-site positions, excluding the theozyme residues. The FuncLib method restricts amino acid mutations to those that are likely to appear in the natural diversity of homologous proteins (*4*). To adapt the design process to the requirements of a *de novo* reaction, we removed all restrictions in the active site. We selected six and twelve designs by system energy for experimental testing for Des61 and Des27 respectively, each comprising 5-8 specific mutations relative to their origin. All designs exhibited high expression yields and showed cooperative denaturation (table S1). One design derived from Des61 showed catalytic efficiency of 3,600 M^-1^s^-1^ and *k*_cat_ of 0.85 s^-1^. Remarkably, eight designs based on Des27 showed increased catalytic activity by 2-100 fold (table S1, S2), with Des27.7, harboring seven mutations relative to Des27, reaching *k*_cat_*/K*_m_ 12,700 M^-1^s^-1^ and *k*_cat_ 2.85 s^-1^, an order of magnitude greater than any previously reported computational design (Figure 2A,B) (*9*). This design diverges significantly from natural IGPSs, and a pairwise sequence alignment to the closest protein in the non-redundant sequence database reveals 141 mutations and multiple insertions and deletions (fig. S3).

**Fig. 2.**
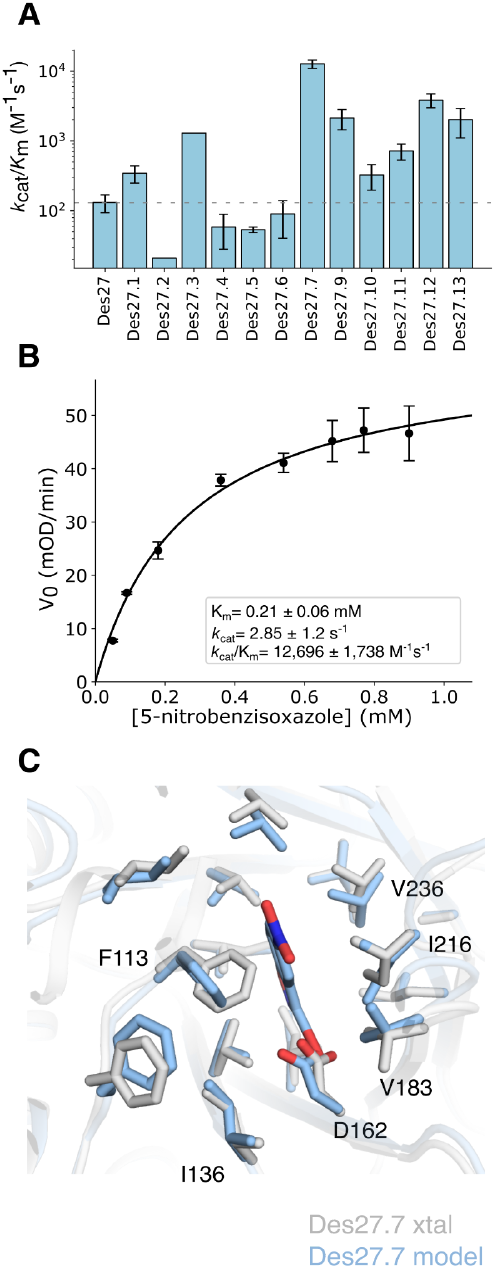
Improving catalytic efficiency through low-throughput screening of FuncLib designs. **A**. Catalytic efficiencies of 12 FuncLib designs encoding 5-8 active-site mutations relative to Des27. **B**. Michaelis-Menten analysis of Des27.7. **C**. The crystal structure of the ligand-unbound Des27.7 (gray, PDB entry 9HVB; to be released upon publication) verifies the accuracy of the designed active site (blue) with rmsd < 0.5 Å. Data are represented as mean ± SD.

We analyzed the structural models of Des27 and its FuncLib-derived variants to understand the mechanistic basis for the three orders of magnitude differences in catalytic efficiency among the designs, using the Rosetta forcefield and molecular dynamics (MD) simulations. A sequence alignment of the FuncLib designs shows that Ile136Val, Ile216Val, and Val183Ile are associated with high catalytic efficiency (Fig. 3A). Contrasting the structure models of Des27 and Des27.7 reveals that these mutations may increase hydrophobic packing around the catalytic Asp162, likely improving its preorganization and desolvation and increasing its reactivity (Fig. 3A, 3B top). Indeed, the Rosetta computed solvation energy of Asp162 is correlated with catalytic efficiency (Spearman ρ=-0.86, *p*-value <10^-3^, Fig. 3A). As further support, MD simulations show that Asp162 is conformationally dynamic in the ligand-unbound models, sampling multiple metastable conformations, and that the fraction of nonproductive conformations decreases in Des27.7 relative to Des27 (fig. S4). Furthermore, the Des27 model suggests that Leu236 may partly overlap with the substrate (Fig. 3B middle), and that the mutation to Val in Des27.7 would alleviate this unfavorable interaction increasing the volume of the pocket from 717 to 829 Å^3^ in Des27.7 relative to Des27 (fig. S5). Finally, Ile54Val, Phe92His and Leu183Val may improve the solvation of the polar nitro moiety of the substrate (Fig. 3B bottom), and Phe92His may enable water-mediated polar interactions with the nitro group. Thus, six of the seven mutations distinguishing Des27 and Des27.7 likely improve the catalytic efficiency by reshaping the active-site pocket for better substrate recognition and optimizing the positioning and reactivity of the catalytic base.

**Fig. 3.**
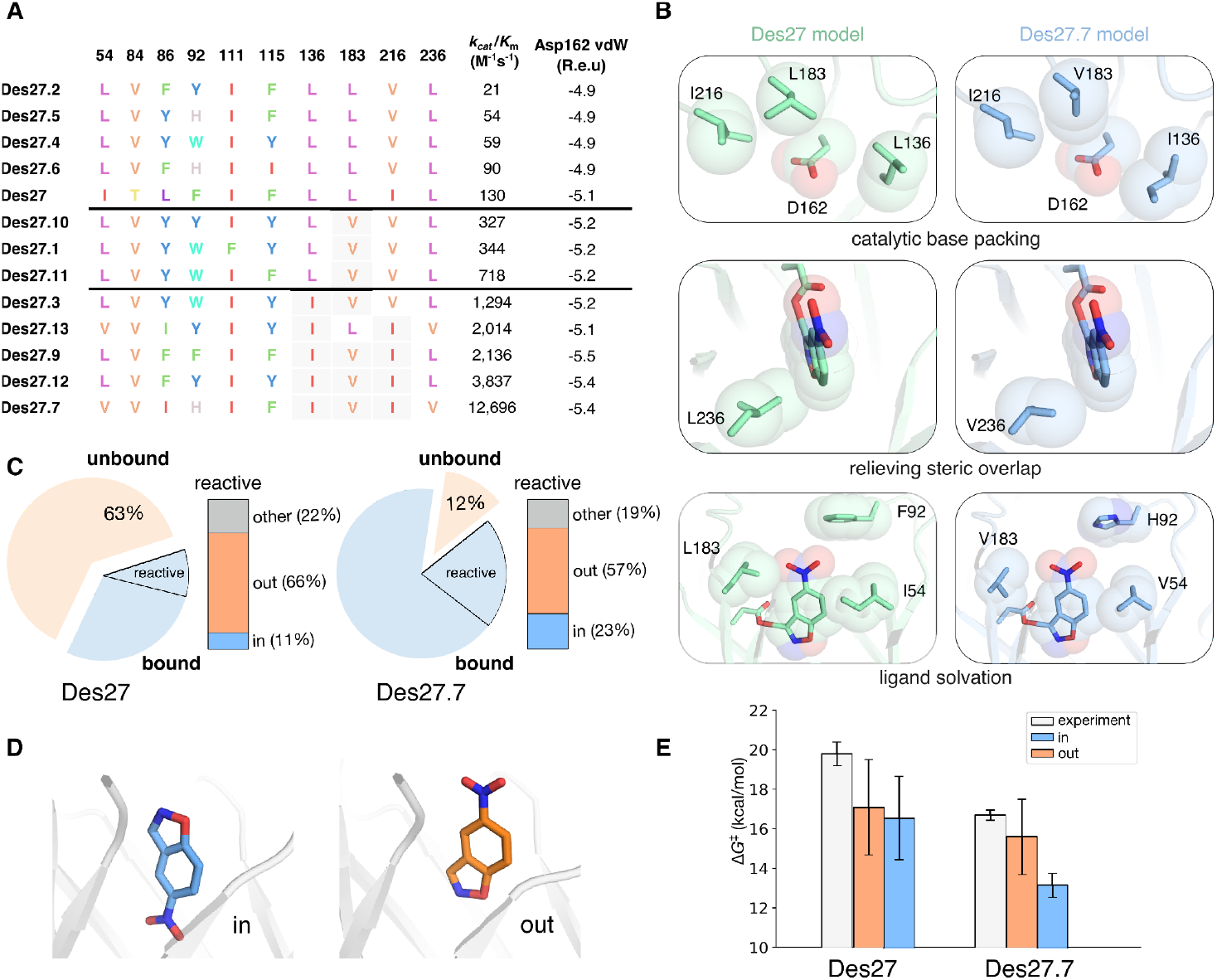
Structural, energetic, and dynamic contributions to the improved catalytic rate of Des27.7. **A**. Mutations (gray background) at positions 136, 216, and 236 trend with increasing catalytic efficiency, which also trends with Rosetta computed van der Waals energy of the catalytic Asp162. R.e.u. – Rosetta energy units. **B**. Analysis of the structural basis of increased KE activity by comparing substrate-bound models of Des27 (left) and Des27.7 (right). **C**. Percentage of MD simulation time in which the substrate is within the active site for Des27 and Des27.7 (<= 4 Å between the substrate and the active site center of mass; blue) or distant from the active site (wheat). 24% of the time spent in the bound conformation the substrate adopts a reactive donor-acceptor geometry (highlighted arc). The right blue bars show the distribution of conformations in the reactive mode. **D**. In MD simulations, 5-nitrobenzisoxazole (sticks) can assume two catalytically competent conformations: one where the nitro group is buried inside the TIM barrel (in, blue) and another in which it is solvent exposed (out, orange). Shown are two representative conformations from the MD simulations. **E**. Activation free energy for the proton abstraction of 5-nitrobenzisoxazole, comparing Δ*G*^‡^ computed from the experimentally determined *k*_cat_ values using the Eyring rate equation assuming *T*=298 K (experiment) and the corresponding values calculated for substrate conformations “in” and “out”.

To analyze the stability of the substrate within the active site, we conducted microsecond MD simulations of Des27 and Des27.7, starting from their ligand-bound design models. In both cases, and across all replicas, the substrate exited and reentered the active-site pocket multiple times (fig. S6), with Des27.7 showing five times more substrate retention (Fig. 3C, table S3). This contrasts with the typical scenario in MD simulations where unbinding events are terminal (*43, 44*), indicating that our designs exhibit improved substrate binding, and that Des27.7 improves it further. We also noticed that the substrate can enter the pocket in two reactive conformations that are inverted: one that closely matches the design model, with the nitro substituent occupying the entrance to the active site, and one in which it is inverted by 180º (Fig. 3D). Empirical valence bond (EVB) calculations of reaction free energies (*45*) show similar energy profiles for both conformations, indicating that both are catalytically competent (Fig. 3E, table S4). Taken together, the MD and EVB calculations suggest that the experimentally measured results reflect the sum of both reaction modes with the “out” conformation (Fig. 3C) being occupied a greater fraction than the “in” conformation but the latter being slightly more reactive (Fig. 3E). Further, while such desolvated nitro group conformations were observed in other *de novo* designed KEs (*12, 43, 44*), in those studies, only one conformation was catalytically competent (*44*). Thus, the high efficiency of Des27.7 may be partly due to the high preorganization of the active-site pocket and its ability to accommodate productive substrate interactions through distinct conformations.

To verify the molecular accuracy of the design process, we determined the structure of Des27.7 in the unbound form by crystallographic analysis (table S5; PDB coordinate file to be released upon publication). All active-site positions aligned well to the design conception (< 0.5 Å root mean square deviation; rmsd), including the catalytic Asp162, although a slight shift (rmsd 0.78 Å) was observed in the orientation of Phe113. Outside the active-site pocket, 180 out of 257 positions aligned with rmsd < 0.6 Å, but 65 amino acids either deviated or did not exhibit significant electron density, likely indicating backbone flexibility in this region (fig. S7). This fragment is known to be dynamic in the IGPS protein family (*46*) but it lies outside the active-site pocket and probably does not contribute directly to reactivity and substrate recognition. Taken together, our results verify a fully computational pipeline that designs an accurate *de novo* active site and generates a stable and high-efficiency new-to-nature enzyme.

### Necessary and sufficient conditions for *de novo* enzyme design

Our computational workflow is based on the combination of several components, each of which designs multiple mutations that address aspects that are critical for efficient biocatalysis, such as backbone diversity, stability, foldability and activity. We next probed whether each of these components contributes to the intended property and whether all are essential.

We started by examining whether modular assembly and design is essential for generating diverse backbones. Instead of applying modular assembly and design, we applied the subsequent steps of the workflow to 1,200 representative IGPSs that were modeled using AlphaFold2 (see Methods). In design round 2 (R2), we tested 55 designs, of which 49 were solubly expressed (89%), and 28 (50%) exhibited apparent cooperative unfolding with *T*_*m*_ values 47-88°C. 70% of the cooperatively folded designs (20 designs) displayed measurable KE activity with *k*_ca*t*_*/K*_m_ in the range of 0.5-155 M^-1^s^-1^, demonstrating that the workflow can design stable and functional KEs in a wide range of different starting points. As expected, designs that did not show cooperative unfolding lacked KE activity. We applied FuncLib to the active sites of six designs and tested 9-14 variants for each starting point. In five cases, catalytic efficiencies improved by 3-10 fold (table S2), with the highest catalytic efficiency reaching 300 M^-1^s^-1^ (R2.Des39.2). We determined the crystallographic structure of two designs, R2.Des39 (*k*_cat_*/K*_m_ 100 M^-1^s^-1^) and Des49 (*k*_cat_*/K*_m_ 150 M^-1^s^-1^) (fig. S8-S9, table S5). The active sites were close to the design conception (rmsd <0.6Å and <0.82Å, respectively), but in both cases, several loops either lacked electron density or exhibited significant conformational changes compared to the designs, which could impede binding (*47*). To explore whether the foldability of these loops could be improved, we applied FuncLib to the design models to stabilize these regions. Three out of 16 FuncLib variants of Des39.2 showed a significant increase in catalytic efficiency, with improvements up to 20 fold compared to the original design, reaching *k*_cat_*/K*_m_ 2,000 M^-1^s^-1^ (table S1, S2). These results demonstrate that AI-based structure prediction of natural enzyme families provides a resource for *de novo* enzyme design, and that the computational workflow reproducibly generates efficient enzymes. In this case, however, the structural diversity afforded by modular assembly and design generates more opportunities for high-efficiency designs.

As a next step to understanding the necessary and sufficient conditions for design of high-efficiency enzymes, we deconvolute the contributions to the high stability and activity of Des27.7 from subsequent design steps. As expected, combinatorial assembly and design alone (with 92 mutations to any natural protein) does not exhibit any KE activity. PROSS stability design (11 mutations) substantially improves both bacterial expression levels and thermal stability (69 °C, compared to 57 °C) (Fig. 4, fig. S10). Implementing the active site from Des27.7 (15 mutations) onto the combinatorial assembly starting point confers high activity levels (2,900 M^-1^s^-1^) but fourfold lower than in Des27.7. We then tested the combination of modular assembly, PROSS and the designed active site, observing a synergistic, higher-than-expected improvement in both stability and reactivity relative to the individual contributions of PROSS and active-site design. Thus, despite the large number of mutations introduced by each computational component, resulting designs do not exhibit the frustrating trade-offs between stability and activity that are often observed in laboratory-evolution campaigns (*2, 13, 48*).

**Fig. 4.**
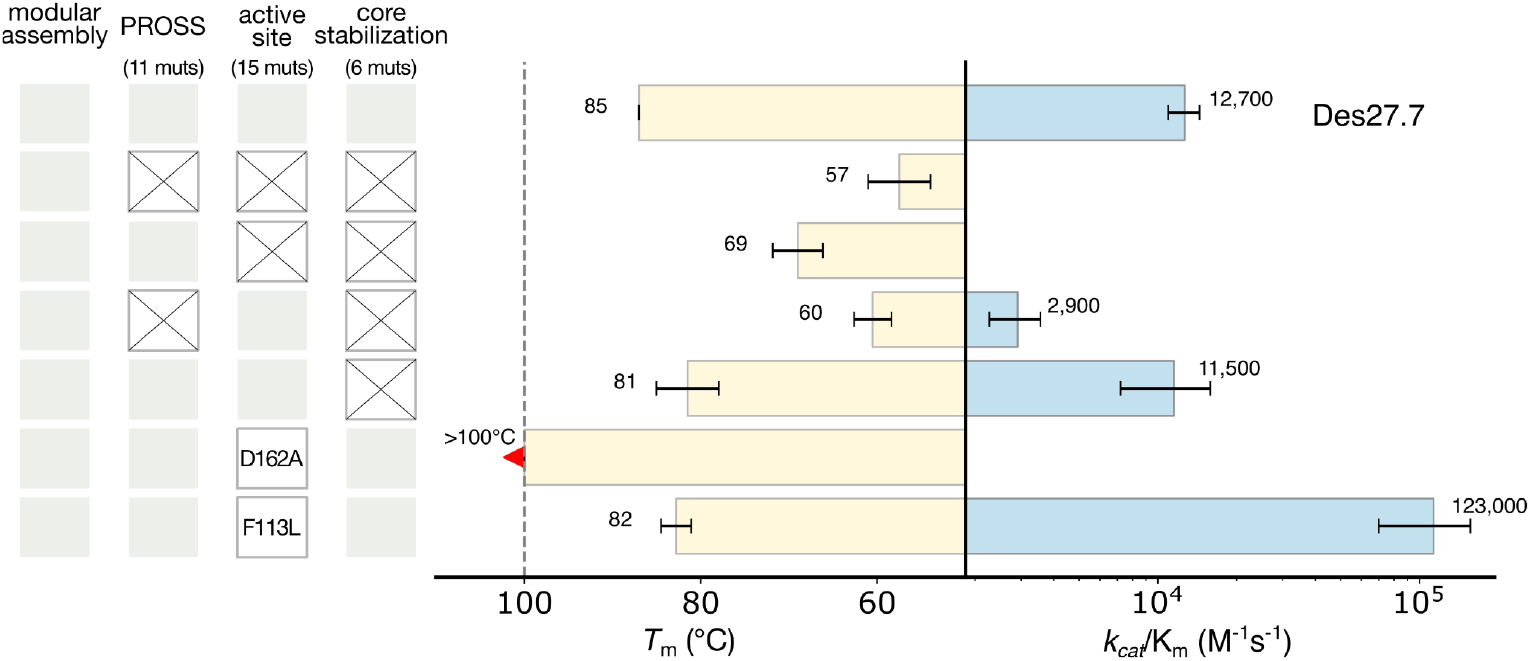
Deconvoluting the contributions of the mutations in Des27.7 reveals necessary and sufficient components for effective *de novo* enzyme design. Apparent melting temperature and catalytic efficiency of Des27.7 (top row) are compared to variants in which design components are ablated (X symbols). The number of mutations relative to the modular assembly is indicated in parentheses. Data are represented as mean ± SD.

Finally, we evaluated the contribution of the theozyme to the activity of Des27.7. Mutating the catalytic base Asp162 to Ala completely abolished activity, verifying that the designed base is essential. Remarkably, this single-point mutation also dramatically increased protein stability, with the apparent melting temperature rising from 85°C in Des27.7 to above boiling point (Fig. 4). This significant increase in stability underscores the strong destabilization induced by desolvating a charged group in the core of the active site and the importance of effective stability design methods.

Additionally, we replaced the second theozyme residue, Phe113, with Leu which was predicted to be tolerated by Rosetta atomistic calculations. Surprisingly, we observed an order of magnitude increase in catalytic efficiency and rate to *k*_cat_/*K*_m_ of 123,000 M^-1^s^-1^ and *k*_cat_ of 30 s^-1^, surpassing by two orders of magnitude recently designed enzymes in AI-generated proteins (*k*_cat_= 0.03-0.7 s^-1^) (*5, 6, 49*). To understand the reasons for this large gain in efficiency, we modeled the Leu mutation in both unbound and transition states. Unlike the reorientation observed for Phe113 between the ligand-bound and unbound structure models of Des27.7 (fig. S11A), Leu113 exhibits almost no sidechain conformation changes (fig. S11B), suggesting improved preorganization. The fact that a completely aliphatic active-site pocket can effectively accelerate the KE reaction is in line with the observation that London dispersion forces are sufficient for transition-state stabilization (*50*). This finding challenges a two decades assumption in computational KE design that an aromatic residue is important for ligand binding (*7, 9*).

## Conclusions

*De novo* enzyme design has so far resulted in rudimentary catalytic rates and required iterative random mutagenesis to close the gap with natural enzymes. Our strategy addresses this challenge by generating diverse TIM barrel backbones, stabilizing the protein, and designing preorganized active-site constellations. This integrative design approach allowed us to explore the principles underlying high stability and activity in KE biocatalysis. In a single step, we generated a dozen designs with activities that spanned three orders of magnitude, offering insights into the determinants of high activity. The best variant showed high stability, and unprecedented catalytic efficiency for a fully designed enzyme (> 85 °C and 12,700 M^-1^s^-1^, respectively), which could be increased to >10^5^ M^-1^s^-1^ with a single point mutation. Active-site preorganization combined with the ability to adopt multiple catalytically competent substrate-binding conformations, distinguishes this design from previously generated ones. Importantly, our best variant exhibited a catalytic rate (30 s^-1^) and efficiency as the median of natural enzymes (*16*). In contrast, prior efforts employing iterative experimental mutagenesis did not achieve such dramatic improvements without testing orders of magnitude more variants. The layered design methodology allowed us to dissect the effects of active-site mutations and stabilizing mutations outside of it, revealing their synergistic impact on activity and stability. Finally, our finding that an aromatic residue is not always optimal for substrate recognition – a decades-long assumption in KE design – underscores the need for a close collaboration between computational chemists and protein designers. We anticipate that the design workflow and this collaboration will generate sophisticated new-to-nature enzymes and deepen our understanding of the fundamentals of biocatalysis.

## Acknowledgments

We thank D. Hilvert for discussions and R. Lipsh-Sokolik, Z. Avizemer, A. Tennenhouse and O. Khersonsky for critical reading. We also thank O. Khersonsky, M. Goldsmith and M. Ovadis for technical help.

