## Supplementary material for "High-efficiency Kemp eliminases by complete computational design": Methods and SI

#### Materials and Methods

Custom Python scripts, RosettaScripts (*51*), command lines, Jupyter notebooks, and datasets used for de novo enzyme design are available at
https://github.com/Fleishman-Lab/denovoKemp

**Backbone generation and stabilization.** Modular assembly and design was applied as described in ref. (*36*). Briefly, five different IGPS structures (PDB entries: 1LBF, 1I4A, 1JCM, 1VC4, 4FB7) were aligned and segmented into five fragments according to points of maximum structure conservation at positions 44, 105, 154 and 206 (numbering relative to PDB entry 1I4N). The fragments were then computationally combined all against all, and Rosetta sequence design was applied to optimize the stability and compatibility between the segments, resulting in 2,500 backbones. Design calculations were constrained using a position-specific scoring matrix (PSSM) that was generated for each structure using PROSS (*3*). For designs 37-73, an additional stabilization protocol was applied. This protocol, based on mutational scanning with PSSM constraints, identifies the most beneficial mutations across the protein. These mutations are combined and threaded onto the input structure (protocol to be published). These backbones were then evaluated by an activity predictor (*35*) and the top 1000 designs were chosen.

To implement the workflow in non-assembled backbones (R2 series), we used AlphaFold2 (*52*) to model the structures of 1,200 IGPSs that exhibited sequence identity between 30-90% to one another. Models with average pLDDT scores <90 were discarded. All structures underwent PROSS stability design calculations (*3*) and Design 8 was selected for further calculations. For AlphaFold2 models, PROSS design was disabled in amino acids that exhibit low predicted accuracy (pLDDT <90%) and 5 Å from these residues.

**Catalytic site generation.** Theozyme geometries (table S6) were based on previous calculations (*7*). Geometric parameters that define the catalytic placements, such as tolerance, penalty coefficient, periodicity, and number of matching samples to test were manually adjusted (*53*). The interaction between the catalytic base and the acidic carbon on the ligand was defined as covalent to mimic transition-state geometry. Theozyme placement was carried out using the Rosetta Matcher algorithm (*37*). All positions inside or in the opening of the active site were allowed for theozyme matching (Fig. 1 step 3).

**Initial active-site design and filtering**. After matching, Rosetta sequence design was performed in an 8 Å shell around the ligand and catalytic residues. The design was performed under theozyme and PSSM constraints. To constrain the sequence space, the catalytic residues of the IGPS family, as described by the M-CSA database (*54*), underwent Rosetta computational mutation scanning, and all mutations with $\Delta\Delta G_{system} < +1$ Rosetta energy units (R.e.u) compared to the starting identity were included as allowed for design. The Rosetta Match and design steps generated $10^5$-$10^6$ designs for each starting structure. Designs were filtered based on a "fuzzy"-logic objective function (*39*) that balanced potentially conflicting criteria: energy density (system energy divided by the protein length), energy rank relative to other designs in the same backbone, active site van der Waals energy (vdW), catalytic base vdW, ligand solvation and accuracy of theozyme geometry. vdW energy is defined as the sum of the Rosetta atomistic energy terms fa_atr and fa_rep (as weighted in Rosetta ref15 scoring function (*55*)).

**Active-site and core stabilization.** To enhance active-site stability, we performed an enumeration of all low-energy mutations in the active site with $\Delta\Delta G_{system} <+3$ R.e.u. and chose the top variant. To ensure amino acid optimality throughout the protein, a pSUFER (*40*) scan was performed on the whole protein excluding the active site. Flagged positions, those with at least five favorable amino acid substitutions ($\Delta\Delta G_{system} < 0$) , were redesigned using FuncLib calculations (*4*). The lowest-energy design was selected.

**Computational validation.** Active-site preorganization was analyzed by performing extensive rigid-body minimization in the absence of the ligand. Structures in which the catalytic base exhibited an rmsd > 1.2 Å relative to the ligand-unbound model were discarded. For the R2 series the workflow included an additional validation step comparing the bound model and the AlphaFold2-predicted model. Designs were accepted if the rmsd between the AlphaFold2 model and the Rosetta model was < 1 Å.

**Active-site optimization.** All functional variants identified through experimental screening were optimized by identifying diverse and stable active-site constellations using FuncLib (*4*). FuncLib employs two filters to constrain the enumerated sequence space: a filter based on homologous sequences and exclusion of destabilizing point mutations. However, in *de novo* design of function, the homologous sequence filter is irrelevant and was omitted. For experimental screening, the 10–15 lowest-energy designs were selected.

**Protein expression.** The designed genes were ordered from Twist Bioscience, cloned into pET28 plasmid with an N-terminal His-tag, followed by a bdSUMO tag. Plasmids were transformed into *E. coli* BL21 (DE3) cells. For expression, 50 ml 2YT medium supplemented with 50 µg/ml kanamycin were inoculated with 500 µl overnight culture produced from a single colony and grown at 37 °C until $OD_{600}$ 0.6-0.8. Overexpression was induced by adding 1 mM IPTG, the cultures were grown for 20 h at 16 °C, harvested, and the pellet was frozen at −20 °C. The cells were resuspended in basic buffer (50 mM Tris-Cl pH 7.25, 200 mM NaCl) supplemented with 10 µg/ml lysozyme, protease-inhibitor cocktail (Sigma) and benzonase, lysed by sonication and centrifuged at 20,000 g for 30 min. The soluble fraction was loaded onto a Ni-NTA (nitrilotriacetic acid) column and washed twice with basic buffer and 20 mM imidazole. The protein was subjected to overnight on-column Sumo protease cleavage at 4 °C (5 µg/ml in basic buffer). Protein purity was assessed by SDS-PAGE. Protein concentration was determined using Pierce BCA protein assay kit. For crystallography, large scale scale-expression was performed in in 1500 ml culture. After Ni-NTA purification and bdSUMO cleavage, the protein was purified by gel filtration (HiLoad 26/600 Superdex75 preparative grade column, GE).

**Activity assay and determination of kinetic parameters.** Product formation was monitored spectrophotometrically at 380 nm in 200-µl reaction volumes using 96-well plates. For initial screening, the reactions were started by adding 150 µl of 1 mM 5-nitrobenzisoxazole in basic buffer to 50 µl of purified protein. 5-nitrobenzisoxazole was used from 0.1 M stock in acetonitrile. For the kinetic characterization, 150 µL of 5-nitrobenzisoxazole at various concentrations (final 0.05–0.75 mM in basic buffer with 1 mM acetonitrile) was mixed with 50 µL purified protein. Kinetic parameters were obtained by fitting the data to the Michaelis–Menten equation $v_0 = k_{cat}[E]_0[S]_0/([S]_0 + K_m)$. At low substrate concentrations the data were fitted to the linear regime of the Michaelis–Menten model $v_0 = [S]_0[E]_0 k_{cat}/K_m$, and $k_{cat}/K_m$ values were inferred from the slope. The reported results are the average of duplicate measurements. All measurements in the main text were performed in biological duplicates or triplicates.

**Thermal stability.** Apparent $T_m$ measurements were performed using nanoscale differential scanning fluorimetry (nanoDSF) experiments (Prometheus NT.Plex instrument, NanoTemper Technologies). The temperature ramp was between 20-95°C with 1.0 °C/min slope.

**Crystallization, Data Collection and Structure Determination**

Crystals were grown at 19°C using the sitting drop vapor diffusion method. AlphaFold2 (*52*) was used to generate all three models for molecular replacement (table S5). Initial models were iteratively rebuilt and refined using COOT (*56*) and PHENIX (*57*). Model geometry was evaluated using MOLPROBITY (*58*). Atomic coordinates and structure factors for Des27.7, R2.Des39, and R2.Des49 are deposited in the PDB database under accession numbers 9HVB, 9HVH, and 9HVG, respectively.

Specific crystallization conditions:

**Des27.7:** The well solution contained 0.15 M lithium sulfate monohydrate, 0.1 M citric acid (pH 3.5), and 18% polyethylene glycol (PEG) 6000. Diffraction data were collected to 2.0 Å resolution at 100 K using an in-house Rigaku liquid-metal-jet (LMJ) X-ray Synergy System with a HyPix Arc 150° detector. Des27.7 crystallized in the $P6_1$ space group, with one subunit in the asymmetric unit.

**R2.Des39:** The well solution contained 0.07 M citric acid, 0.03 M Bis-Tris propane (pH 3.4), and 14% PEG 3350. Diffraction data were collected to 2.1 Å resolution. R2.Des39 crystallized in the $P2_1$ space group, with two subunits in the asymmetric unit.

**R2.Des49:** The well solution contained 8% Tacsimate (pH 7.0) and 20% PEG 3350. Diffraction data were collected to 1.9 Å resolution. R2.Des49 crystallized in the $C222_1$ space group, with one subunit in the asymmetric unit.

**Molecular Dynamics (MD) and empirical valence bond (EVB) simulations**

**System Setup.** Two designed Kemp eliminases, Des27 and Des27.7, were simulated in both their ligand-bound and unbound forms, using molecular dynamics (MD) simulations to model dynamics, and empirical valence bond (EVB) simulations (*45*). All simulations were initiated from the FuncLib design models for each variant. Ligand-bound simulations were performed in complex with the substrate 5-nitrobenzisoxazole. The partial charges for the substrate were calculated using restrained electrostatic potential (RESP) (*59*) fitting at the HF/6-31G(d) level of theory with Antechamber (*60*), based on gas-phase geometries optimized at the B3LYP/6-31G(d) level of theory in Gaussian 16 Rev. B.01 (*61*). All other force field parameters for the substrate were obtained from the General AMBER Force Field (GAFF2) (*62*). Residue protonation states were checked using PROPKA 3.0 (*63*, *64*) to estimate sidechain p$K_a$s, coupled with visual examination using PyMOL, based on which all residues were kept in their standard protonation states at physiological pH. For simulations of the unbound system, the catalytic residue Asp162 was modeled in its protonated form. All systems were solvated in a truncated octahedral water box containing OPC water molecules (*65*), extending 11.0 Å from the protein in all directions. Neutralization was achieved using 12 $Mg^{2+}$ and 12 $Cl^-$ counterions for the three variants. Protonation patterns of histidine residues for each system are collected in table S7. Non-standard substrate parameters are provided in table S8 and in the Zenodo data package available at DOI: 10.5281/zenodo.14563437.

**Classical Molecular Dynamics Simulations.** All MD simulations in this work were performed using the HIP-accelerated version of Amber24 (*66*) using the ff19SB force field (*67*) and the OPC water model (*65*). MD simulations for all systems followed the same protocol, used also in prior work modeling designed Kemp eliminases (*44*). For a detailed description, see ref. (*44*). In brief, each trajectory was first energy minimized with 100 steps of the steepest-descent algorithm, followed by 900 steps of conjugate gradient minimization, applying a 100 kcal $mol^{-1}$ $Å^{-2}$ restraint to all solute (protein and substrate) atoms. The system was then heated from 50 to 300 K in an NVT ensemble using simulated annealing, reaching 300 K within the first 100 ps and continuing for a total of 1 ns with a 1 fs time step. Langevin temperature control (*68*) was employed with a collision frequency of 1 $ps^{-1}$. During this stage, the 100 kcal mol$^{-1}$ Å$^{-2}$ solute restraints were maintained and subsequently reduced to 10 kcal mol$^{-1}$ Å$^{-2}$ in later equilibration steps. A second energy minimization and heating step followed, with positional restraints applied to solute heavy atoms. During subsequent equilibration, the restraints were progressively reduced from 10 to 1 to 0.1 kcal mol$^{-1}$ Å$^{-2}$ before being fully removed. The systems, now with only distance restraints applied, underwent final equilibration for 1 ns in an NPT ensemble (300 K, 1 atm) using a Berendsen barostat (*69*) with a 1 ps pressure relaxation time and Langevin temperature control (collision frequency of 1 ps$^{-1}$). The SHAKE algorithm (*70*) was applied to constrain all bonds involving hydrogen atoms, and all equilibration simulations used a 1 fs time step. Production MD runs were performed using a 4 fs time step, enabled by hydrogen mass repartitioning (*71*) and the SHAKE algorithm (*70*) with an 8 Å direct space nonbonded cutoff, Langevin temperature control (collision frequency of 1 ps$^{-1}$), and a Berendsen barostat (pressure relaxation time of 1 ps). Equilibration of these trajectories is shown in Figure S12. The final production trajectories were 1 μs long for each system, with 5 independent replicas per system, resulting in a total of 5 μs of simulation time per system and 15 μs across all systems.

**Empirical Valence Bond Simulations.** Empirical valence bond simulations were performed on the Des27 and Des27.7 variants, using both substrate conformers (in and out, Fig. 3C) observed as being dominant in the MD simulations, and following the same protocol described in detail in ref. (*44*). Prior to the EVB simulations, in all cases, the enzyme-substrate complex was minimized with Amber24 (*66*) in vacuum, with 2500 steps of the steepest-descent algorithm, followed by 2500 steps of conjugate gradient minimization, applying 10 kcal mol$^{-1}$ Å$^{-2}$ positional restraints on all heavy (non-hydrogen) atoms. The minimization was repeated with the same steps, with 5 kcal mol$^{-1}$ Å$^{-2}$ positional restraint on protein $C_\alpha$-atoms, and substrate heavy atoms, and with twice as many steps, keeping the restraint only on the substrate heavy atoms.

All EVB simulations were performed using the *Q*6 simulation package (*72*), the OPLS-AA force field (*73*), the TIP3P water model (*74*) and the surface constrained all atom solvent (SCAAS) model (*75*) to describe solvent. Long-range interactions were described using the local reaction field (LRF) approach (*76*). Protonation states of ionizable residues within the explicit simulation sphere, as well as histidine protonation patterns (both of which were validated by PROPKA 3.0 (*63*, *64*) and visual inspection), can be found in table S9. Each system was simulated in 30 replicas of 30 ns equilibration, with 5 kcal mol$^{-1}$ Å$^{-2}$ distance-based restraints applied between the substrate hydrogen donor carbon and the acceptor oxygen of Asp162. Each equilibration was followed by 10.2 ns of EVB simulations (200 ps window over 51 discrete EVB windows), carried out without the distance restraint applied, leading to a cumulative 612 ns of EVB simulation time per system (including "in" and "out" substrate conformations), and 3.6 μs of EVB equilibration time and 1.2 μs of EVB simulation time across all systems studied in this work (4.8 μs simulation time in total). The corresponding root mean square deviations of the equilibration phase (calculated with the *Q*Calc6 module of *Q*6) are shown in fig. S13. Representative stationary points for the Kemp elimination reaction catalyzed by Des27, extracted from EVB simulations of this system, are shown in fig. S14. To extract the conformations representing each stationary point, all 30 replicas were evaluated together. The MDTraj software (*77*) was applied to convert the trajectories to a CPPTRAJ compatible format, and clustering was performed based on the RMSD of the substrate heavy atoms, using the average linkage clustering method, with an ε value of 0.75. We note that the key stationary points for Des27.7 are visually similar to those for Des27. Sample input files, parameter files, starting structures and simulation snapshots, have all been submitted to Zenodo, DOI: 10.5281/zenodo.14563437.

**Simulation Analysis.** Unless otherwise stated, all MD analyses were performed using the CPPTRAJ module (*78*) of AmberTools24 (*79*). Trajectory frames were extracted every 400 ps, and results (where applicable) are reported as averages and standard deviations over 5 × 1 μs trajectories per system. The fraction of unbound and bound modes during the simulations were determined by counting trajectory frames. A bound mode was defined based on the distance between the ligand and the center of mass of the active site, including residues 54, 84, 86, 92, 136, 162, 183 and 236. A threshold of 4Å was defined to classify the frames into unbound or bound. Unbound modes have left the active site pocket, but not necessarily dissociated from the protein itself (sampling non-productive conformations out of the active site). The substrate orientation was defined using the distance between the $C_{\alpha}$-atom of residue Leu41 and the N1 and N2 atoms of the substrate. Conformations with a Leu41-N1 distance between 0 and 15 Å and L41-N2 distance between 15 and 20 Å were classified as "out", and otherwise as "in". Pocket volumes of Des27 and Des27.7 systems were calculated using MDPocket (*80*) (*81*), with snapshots taken every 4 ns of the simulations for unbound systems. Additionally, the volume of the ligand was calculated using the mol_volume package in VMD (*82*). Finally, PyMOL was used for all visualization analysis.

### Supplementary Figures

A.

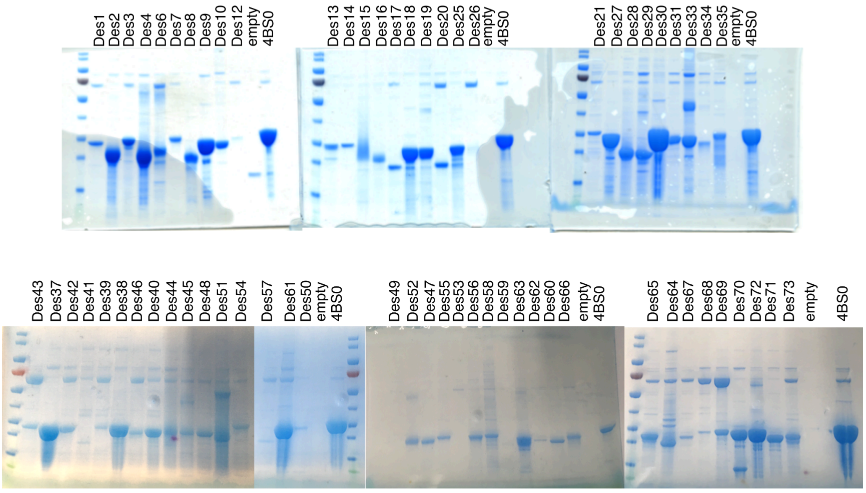

B

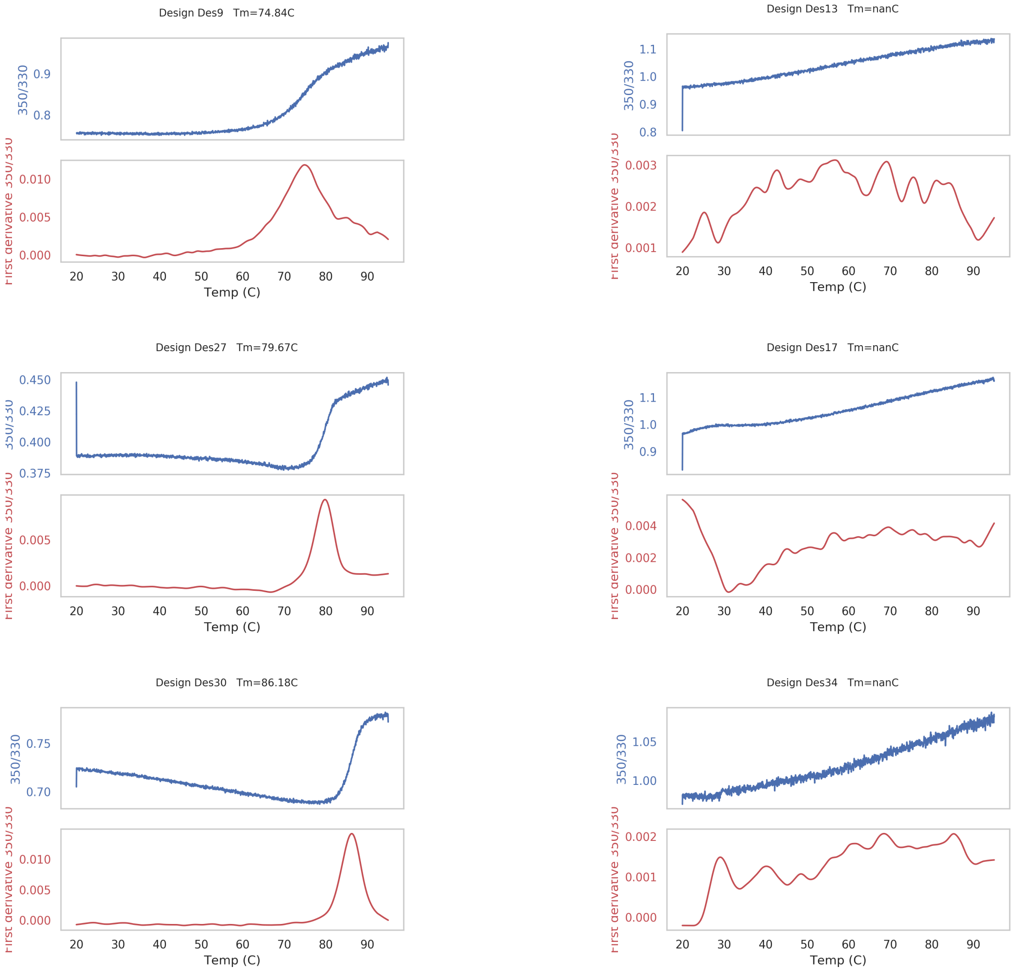

**Fig. S1. Expression and stability of initial design round**. A. 66 designs were solubly expressed. B. Temperature melts of representative cooperatively folded (left) and unfolded (right) designs.

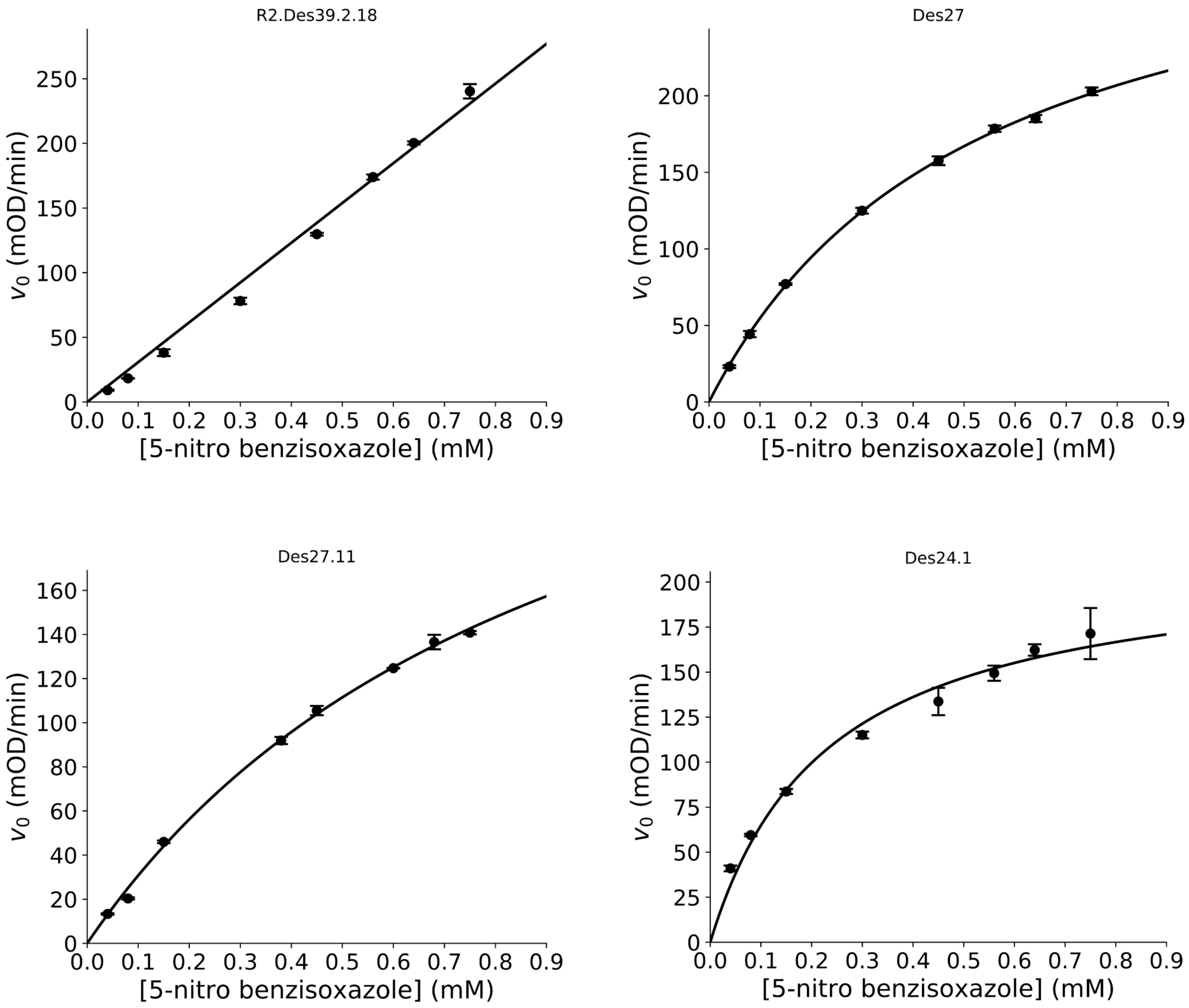

**Fig. S2. Representative Michaelis-Menten plots.** The data were fitted to the Michaelis-Menten equation $v_0=k_{cat}[E]_0[S]_0/([S]_0+K_m)$. When substrate saturation could not be achieved due to limited substrate solubility, the data were fitted to the linear region of the Michaelis-Menten model $v_0=[S]_0[E]_0k_{cat}/K_m$, and $k_{cat}/K_m$ was deduced from the slope. Error bars represent the SD of technical duplicates.

| Score | Expect | Method | Identities | Positives | Gaps |
|---|---|---|---|---|---|
| 196 bits(499) | 7e-58 | Compositional matrix adjust. | 118/259(46%) | 168/259(64%) | 7/259(2%) |

```
Query  2    PSALDAIVADVREDVAAREAVVPFDEIKERAARAPPPRDVLAALRAPGVGIIAVYLRKSP
Sbjct  10   ..V..S..DG....L.V...A.D.....R...K.A..H..M.T.....I.V..EVK.R..

Query  62   SGLDVE--RDPIEYAKT-AEKYAVALVVITDEKYHNGSYEDLEKIRSAVDIPVICFDFIV
Sbjct  70   .KG.LATIS..A.L.ASY..GG.RVIS.L.EQRRFH..LA..DAV.A.....ILRK....

Query  119  DPYQIYLARAYQADAIVLILSVLDDEQYRQLAAVAHSLNMGVIVDVHTEEELERALKAGA
Sbjct  130  S...VHE...HG..LVL..VAA.EQNVLVA.LDRVE..G.TAL.E......AD...E...

Query  179  EIIGIVNQDLKTFEVDRNTAERLGRLARERGFTGVLLAIGGYSTKEELKSMRGL-FDAVV
Sbjct  190  GL..VNARN.H.L..N.SI---F.QI.PGLPNDVLRV.ES.VRGPGD.LTYA.WGA...L

Query  238  IGESLMRAPDPEKAIRELV  256
Sbjct  247  V..G.VTSG..QS.V.S..  265
```

**Fig. S3. Protein BLAST search with Des27.7 sequence as query against the nr database.** Top hit in nr. Mutations are in red, hyphens indicate insertions and deletions.

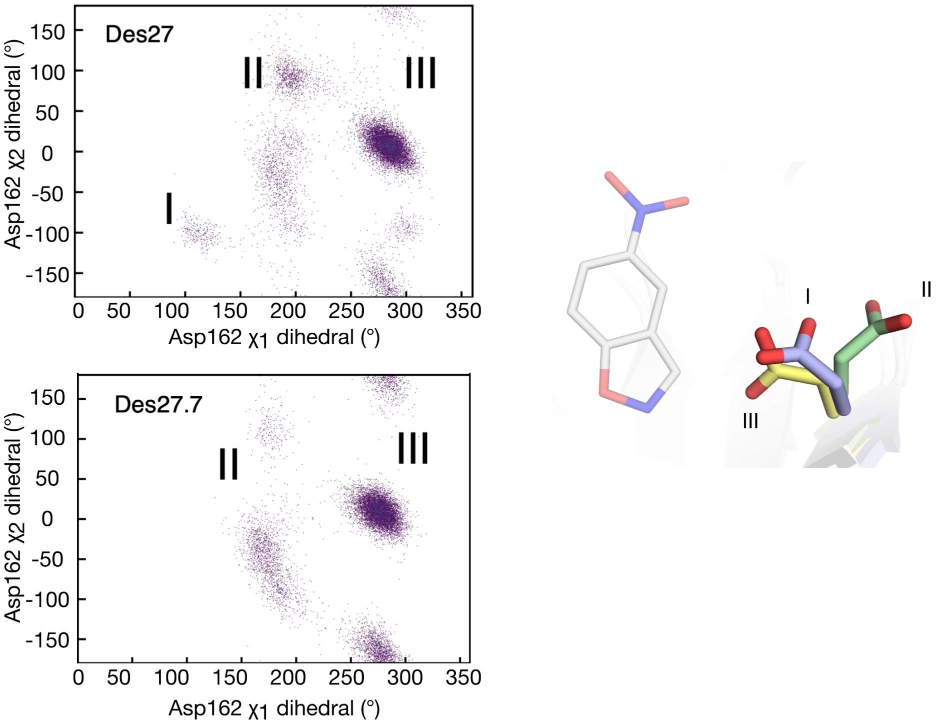

**Fig. S4. Joint distribution of the Asp162 conformational space in the unbound state**. Conformation sampled by the $\chi_1$ and $\chi_2$ dihedral angles of the Asp162 side chain along MD simulations of the Des27 and Des27.7. As can be seen from this data, in the Des27 variant, we observe three distinct metastable Asp162 conformations, which are illustrated in the right panel. Conformation numbers correspond to stick representations on the right, with 5-nitrobenzisoxazole depicted in white sticks for reference.

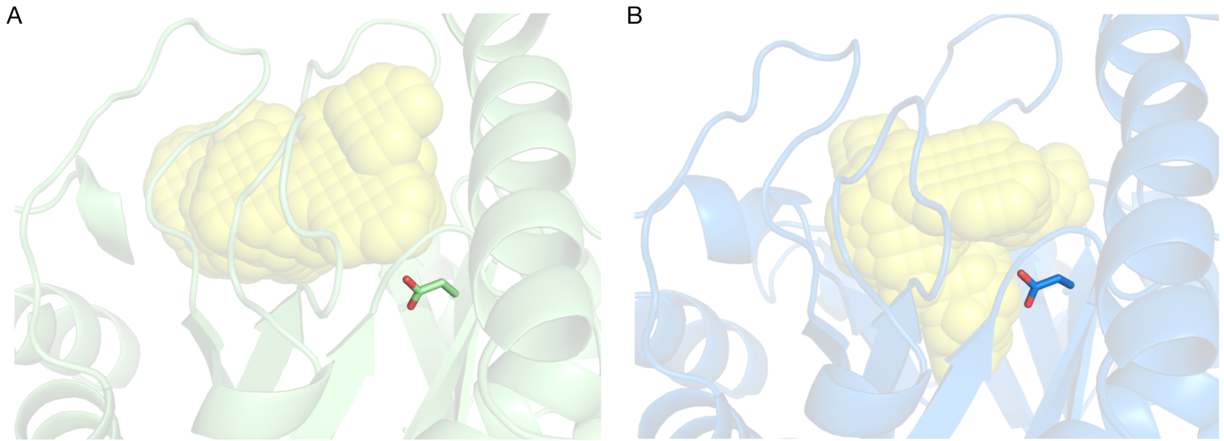

**Fig. S5. Visual representation of the calculated pockets using MDPocket isosurfaces.** Yellow spheres represent pocket volume. Asp162 in sticks. A) Des27 and B) Des27.7. Des27.7 exhibits an active site pocket that can better accommodate 5-nitrobenzisoxazole.

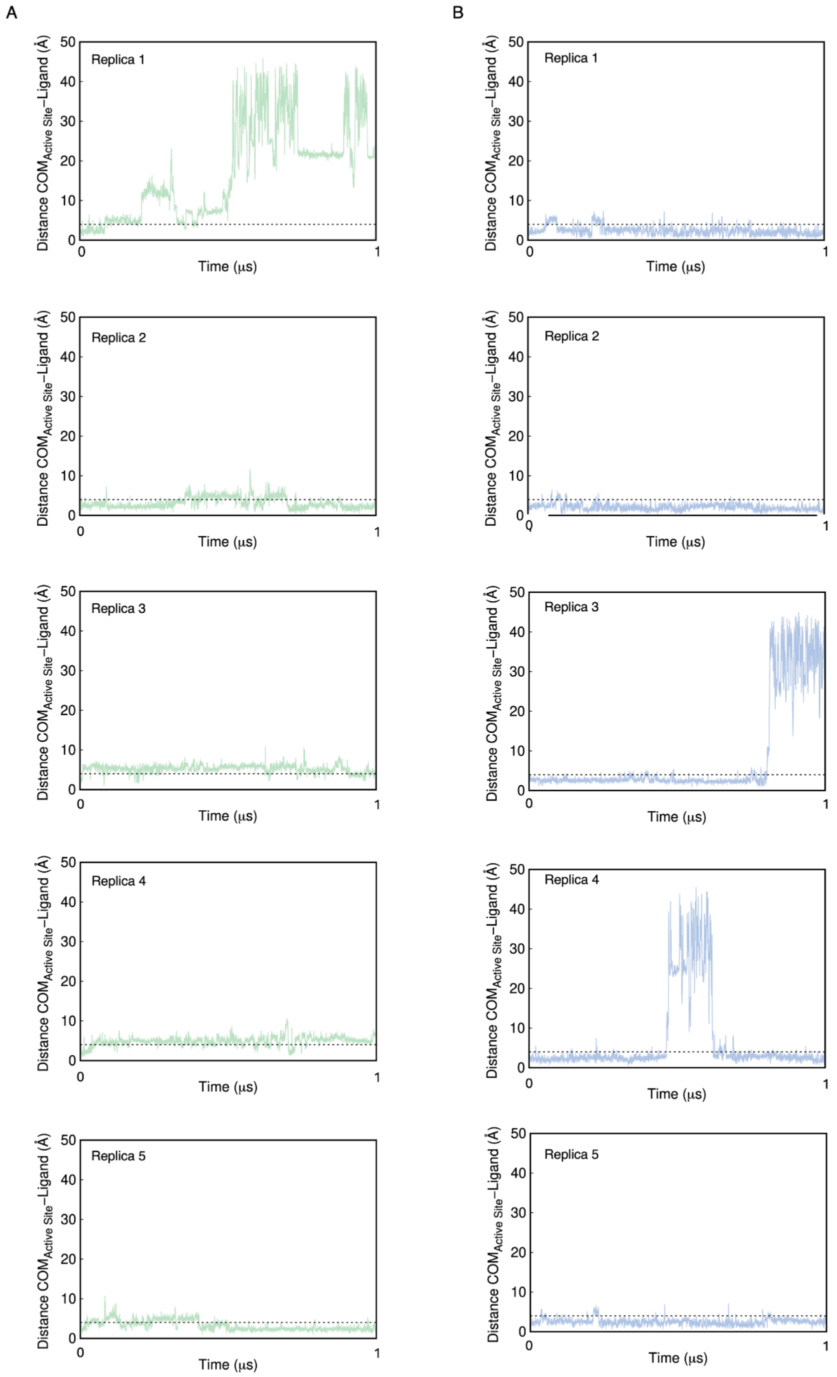

**Fig. S6. Time evolution of the active site center of mass (COM) and ligand distance during total 5 x 1 μs of MD simulations.** A) Des27and B) Des27.7. The dotted gray line at 4 Å marks the threshold for defining the ligand as being within the active site. In two out of ten replicas across the two systems, we observe ligand dissociation without rebinding towards the end of the trajectory on the timescale of our simulations; in a third, in the case of Des27.7, we observe substantive dissociation mid-trajectory with ligand rebinding within our simulation timeframes.

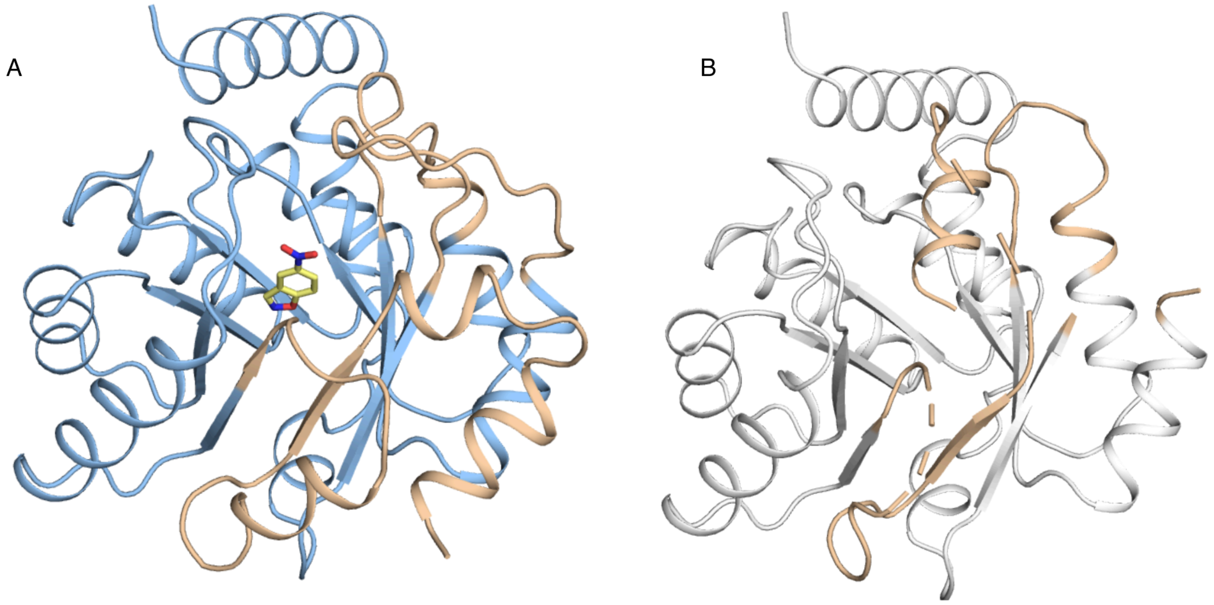

**Fig. S7. Model and crystallographic structure of Des27.7**. A) Des27.7 model. The substrate, 5-nitrobenzisoxazole, in yellow stick. Loop regions in wheat represented misfolded/missing regions in the experimentally determined structure. B) Des27.7 crystallographic structure (PDB 9HVB to be released upon publication). Missing density and misfolded loop in wheat.

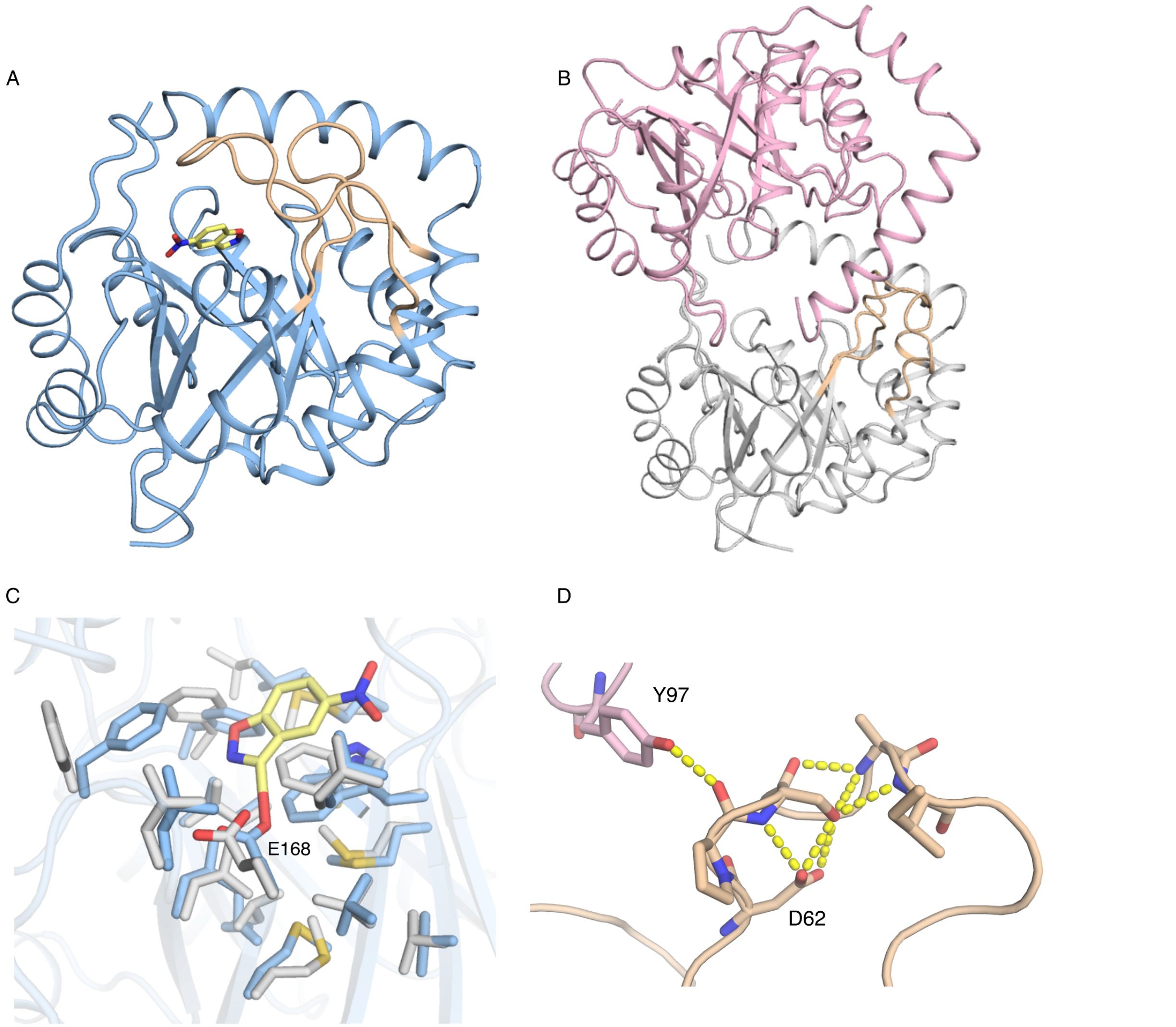

**Fig. S8. Model and crystallographic structure of R2.Des39.** The substrate, 5-nitrobenzisoxazole, is shown in yellow sticks. A) R2.Des39 model. Loop regions in wheat represent misfolded/missing regions in the crystallographic structure. B) R2.Des39 crystallographic structure (PDB 9HVH to be released upon publication). R2.Des39 crystallized as a dimer in the asymmetric unit with few crystallographic contacts that stabilize the wheat-colored loop. The two monomers are in pink and white. Wheat-colored loop corresponds to the one in panel A. C) Active site model (blue sticks) *vs* structure (white sticks). rmsd between the active sites is 0.61 Å. Catalytic Glu168 fits the modeled rotamer and there are only subtle rotameric changes in other active-site residues. These changes do not clash with the modeled ligand. D) Asp62 stabilizes an alternative conformation in the crystal structure. Tyr97 (pink sticks) is from the second monomer. Colors as in panel B.

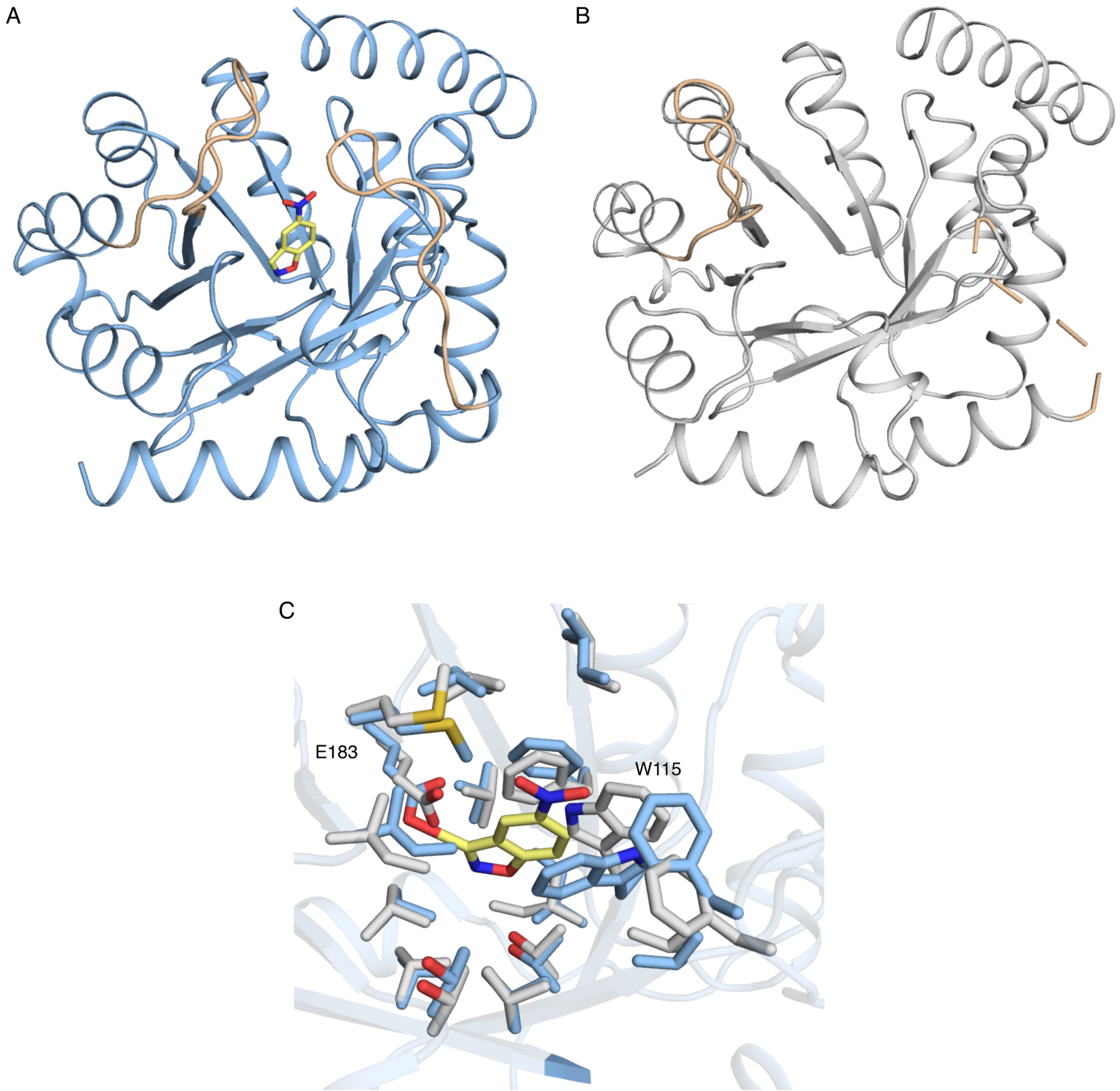

**Fig. S9. Model and crystallographic structure of R2.Des49**. Substrate, 5-nitrobenzisoxazole, is shown in yellow sticks. A) R2.Des49 model. Loop regions in wheat represent misfolded/missing regions in the crystallographic structure. B) R2.Des49 crystallographic structure (PDB 9HVG to be released upon publication). Missing density and misfolded loop in wheat color. C) Active-site model (blue sticks) *vs* structure (white sticks). rmsd between the active sites is 0.82 Å. Trp115 acquires a rotamer different than modeled and might overlap with the ligand.

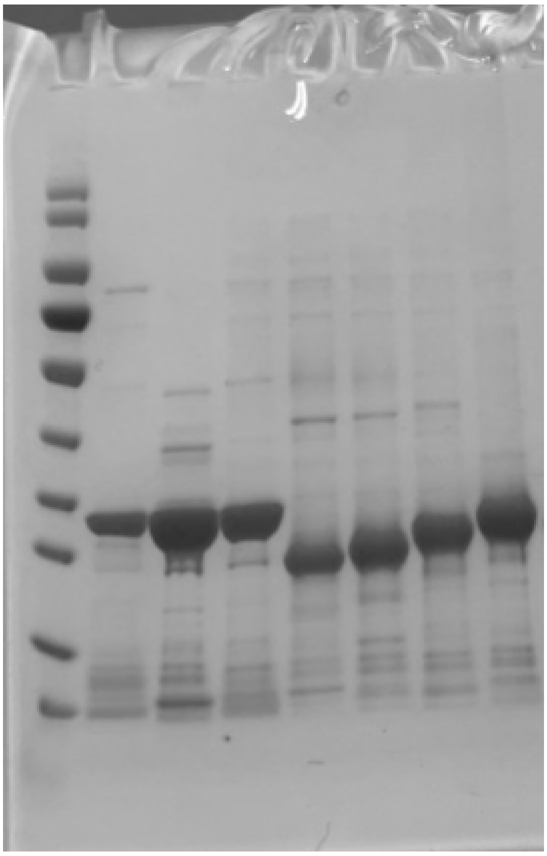

**Fig. S10. Protein expression of designs from ablation study of Des27.7 (relative to main Figure 3).** From left to right: marker, modular assembly, PROSS, modular assembly with active site, Des27.7 Asp162Met, Des27.7 Asp162Ala, Des27.7 Phe113Leu, Des27.7 without core stabilization.

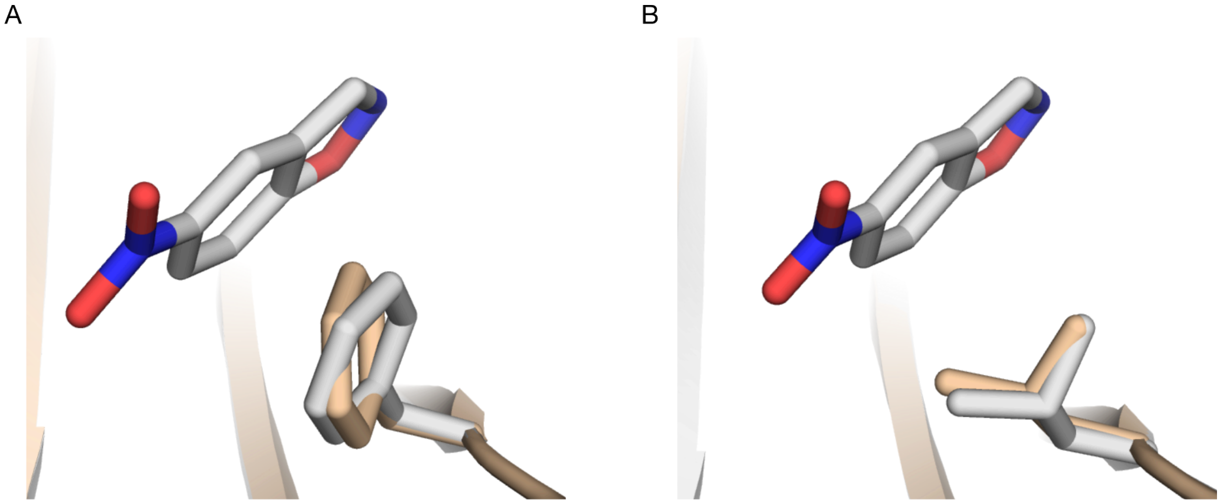

**Fig. S11. Comparison of substrate-bound and unbound structure models of Des27.7 Phe113Leu.** A) A slight sidechain conformational change is observed, with an rmsd of 0.28 Å between the substrate bound and unbound models of Des27.7 Phe113. B) No sidechain shift is observed in the case of Des27.7 Leu113. White sticks represent the substrate bound state, wheat sticks represent the unbound state.

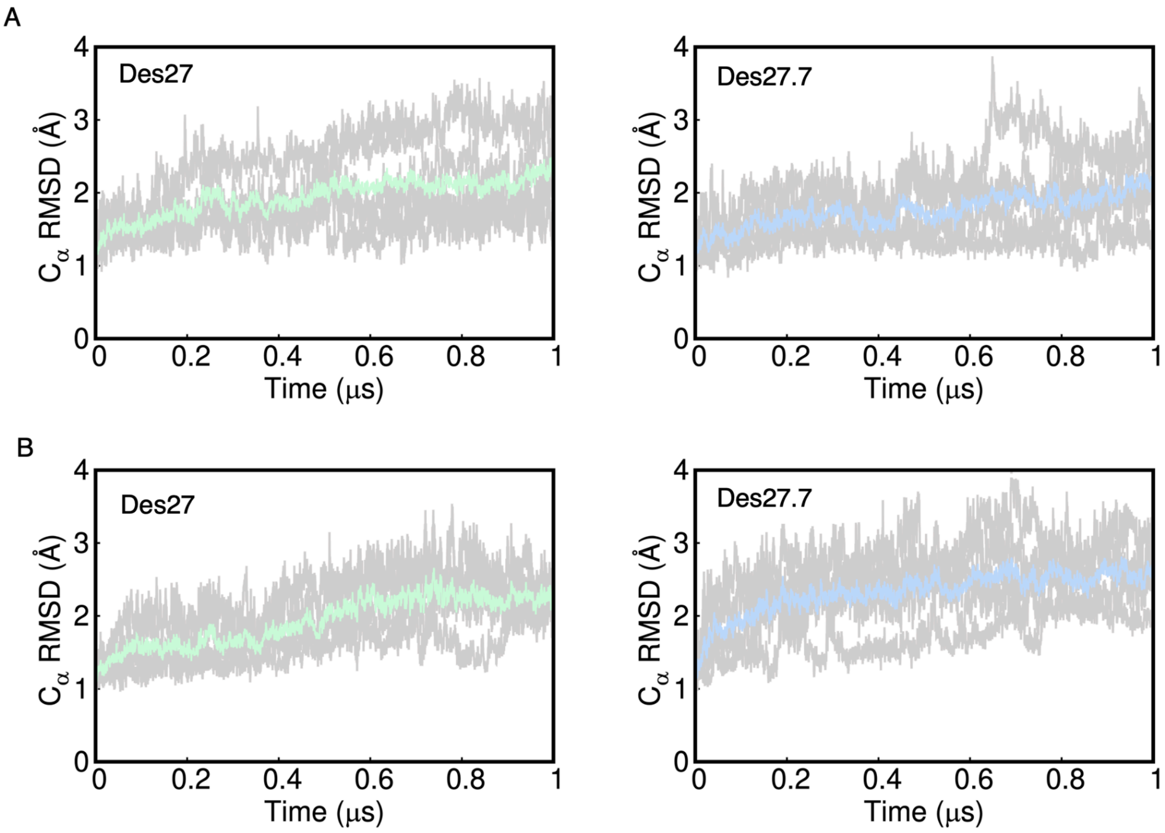

**Fig. S12. Root mean square deviations (rmsd, Å) of the $C_{\alpha}$-atoms from MD simulations.** A) rmsd for unbounded systems Des27 (left), Des27.7 (right) B) rmsd for bounded systems Des27 (left), Des27.7 (right). Data was collected every 400 ps from 5 replicas of 1 μs length each. The gray lines show the five individual runs, and the colored solid line shows a rolling average of the rmsd from all five replicas for each system.

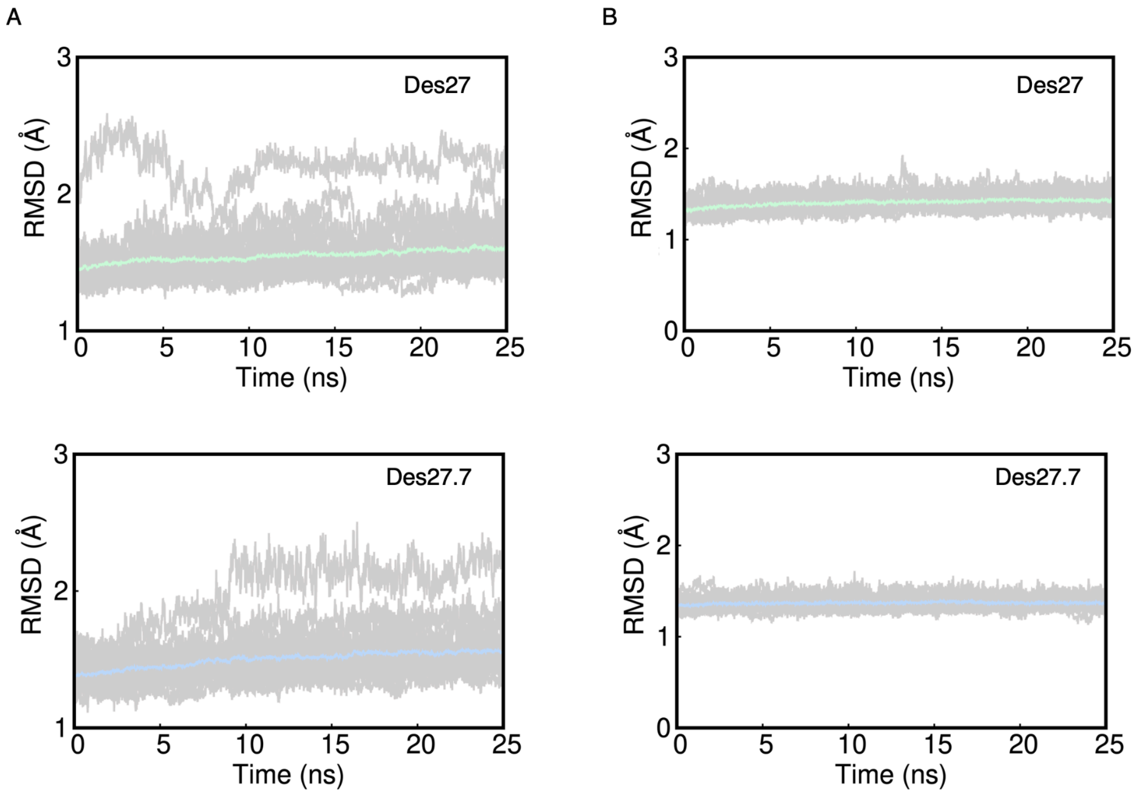

**Fig. S13 Root mean square deviations (rmsd, Å) of all solute atoms of the Des27 and Des27.7 calculated for the equilibration phase prior to our EVB simulations.** A) rmsd for EVB equilibration from "out" substrate conformation. B) rmsd for EVB equilibration from "in" substrate conformation. Data were collected every 10 ps from the initial equilibration runs and shown as averages and standard deviations over ten individual 25 ns MD simulations per system (*i.e.,* 750 ns cumulative simulation time per system). The average rmsd per system is denoted by the colored solid line, and the standard deviations per point over all trajectories are illustrated by the shaded area on each plot.

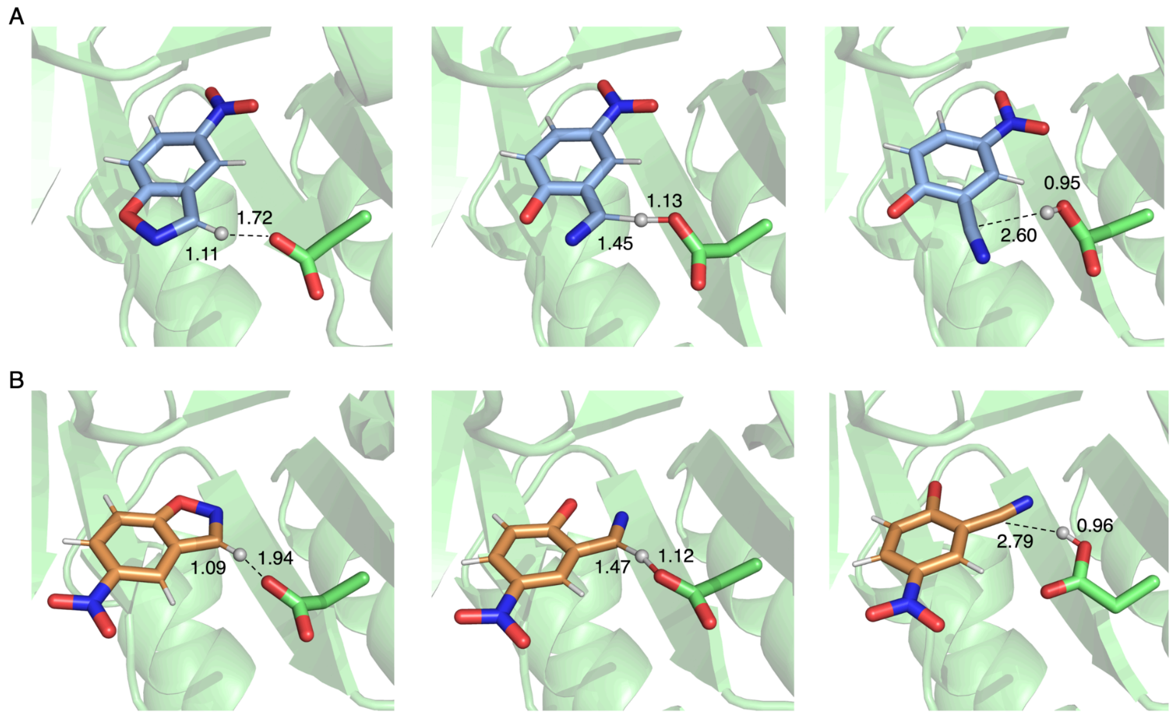

**Fig. S14. EVB Representative structures of the Des27 system**. Michaelis complex (MC, left panel), transition state (TS, middle panel), and product complex (PC, right panel) for the Kemp elimination reaction catalyzed by this enzyme, extracted from EVB trajectories of this reaction. A) For "in" ligand conformation. B) For "out" ligand conformation. Structures were selected based on clustering analysis. The clustering was performed at the MC, TS and PC independently, in order to obtain representative structures for each state. Donor-acceptor distances (Å) are shown for each stationary point. These values are averages of the snapshots taken every 5 ps of the trajectory, determined based on the combined evaluation of 30 replicas.

**Supplementary Tables:**

**Table S1.** Apparent thermal stability and catalytic parameters of designed KEs

| | $k_{cat}$ ($s^{-1}$) | $K_m$ (mM) | $k_{cat}/K_m$ ($M^{-1}s^{-1}$) | $T_m$ (℃) |
|---|---|---|---|---|
| Des27 | 0.07 ± 0.02 | 0.5 ± 0.05 | 131 ± 37 | 79 |
| Des27.1 | n.c. | n.c. | 344 ± 94 | 73 |
| Des27.2 | n.c. | n.c. | 21 | 83 |
| Des27.3 | 1.40 | 1.00 | 1,294 | 81 |
| Des27.4 | 0.03 | 0.78 | 59 ± 30 | 82 |
| Des27.5 | n.c. | n.c. | 54 ± 4 | 83 |
| Des27.6 | n.c. | n.c. | 90 ± 49 | 84 |
| Des27.7 | 2.85 ± 1.20 | 0.22 ± 0.06 | 12,696 ± 1738 | 85 |
| Des27.9 | 3.10 ± 0.14 | 1.55 ± 0.64 | 2,136 ± 692 | 82 |
| Des27.10 | n.c. | n.c. | 327 ± 129 | 81 |
| Des27.11 | 0.64 ± 0.23 | 0.89 ± 0.09 | 718 ± 183 | 80 |
| Des27.12 | 5.15 ± 2.47 | 1.30 ± 0.28 | 3,837 ± 864 | 80 |
| Des27.13 | 2.45 ± 1.48 | 1.15 ± 0.21 | 2,014 ± 907 | 86 |
| De61 | 0.3 | 1.3 | 213 | 65 |
| Des61.1 | 0.85 ± 0.02 | 0.27 ± 0.06 | 3,205 ± 555 | 63 |
| Des61.2 | 0.65 | 0.48 | 1,347 | 58 |
| Des61.3 | 0.6 | 0.76 | 770 | 70 |
| Des61.4 | n.d. | n.d. | n.d. | 67 |
| Des61.5 | n.c. | n.c. | 542 | 62 |
| Des61.6 | 0.77 | 0.43 | 1,778 | 58 |
| R2.Des39 | n.c. | n.c. | 92 | 69 |
| R2.Des39.2 | n.c. | n.c. | 298 ± 58 | 81 |
| R2.Des39.2.1 | n.c. | n.c. | 1,136 | n.m. |
| R2.Des39.2.17 | n.c. | n.c. | 1,121 | n.m. |
| R2.Des39.2.18 | n.c. | n.c. | 2,044 ± 163 | n.m. |
| MA | n.d. | n.d. | n.d. | 60 |
| MA + PROSS | n.d. | n.d. | n.d. | 71 |
| MA + active site | 2.4 ± 1 | 0.80 ± 0.23 | 2,911 ± 642 | 62 |
| MA+ PROSS + active site | 3.8 ± 1 | 0.34 ± 0.06 | 11,531 ± 4,323 | 79 |
| Des27.7 D162A | n.d. | n.d. | n.d. | >100 |
| Des27.7 F113L | 30.0 ± 8 | 0.25 ± 0.03 | 123,274 ± 40,707 | 84 |

* n.c. not calculable;  n.d. not detectable; n.m. not measured; MA-modular assembly

**Table S2.** Sequences and experimental summary of Round1 and Round2 *de novo* designs and selected variants for optimization. (separate file)

**Table S3.** Percentage of simulation time that substrate spends in unbound, bound and reactive modes for Des27 and Des27.7.

| | Unbound | Bound | Reactive[a] | in[b] | out[b] | other[b] |
|---|---|---|---|---|---|---|
| Des27 | 63% | 37% | 9% | 11% | 66% | 23% |
| Des27.7 | 12% | 88% | 21% | 24% | 57% | 19% |

[a] Percentage calculated relative to the total.
[b] Percentage calculated relative to the reactive proportion.

**Table S4.** Experimental and calculated activation free energies for the Kemp elimination of 5-nitrobenzisoxazole by Des27 and Des27.7 design models.[a]

| | Experimental | in | out |
|---|---|---|---|
| Des27 | 19.8 ± 0.6 | 16.6 ± 2.1 | 17.1 ± 2.4 |
| Des27.7 | 16.7 ± 0.3 | 13.2 ± 0.6 | 15.6 ± 1.9 |

[a] All energies are shown in kcal $mol^{-1}$. Experimental activation free energies were obtained from $k_{cat}$ values presented in Table S1, using transition state theory. Calculated values are average values and standard deviations over 30 independent empirical valence bond (EVB) simulations per system, as described in the Material and Methods section. “in” and “out” substrate conformations denote energies obtained from simulations initiated from the substrate starting in the corresponding in/out conformation.

**Table S5.** Crystallographic data and refinement statistics for Des27.7, R2.Des39 and R2.Des49. PDB coordinate files to be released upon acceptance.

| | **Des27.7** | **R2.Des39** | **R2.Des49** |
|---|---|---|---|
| *Data collection* | | | |
| PDB code | 9HVB | 9HVH | 9HVG |
| Wavelength (Å) | 1.34 | 1.34 | 1.34 |
| Resolution range (Å) | 25.23-2.0 (2.07-2.00) | 21.24-2.1(2.17-2.10) | 21.06-1.9(1.97-1.90) |
| Space group | $P\ 6_1$ | $P\ 2_1$ | $C\ 2\ 2\ 2_1$ |
| Unit Cell (Å) | 98.54 98.54 40.48 | 48.84 69.55 79.60 | 48.53 88.94 131.18 |
| (°) | 90 90 120 | 90 101.04 90 | 90 90 90 |
| Unique reflections | 15,405 (1,519) | 30,649 (3,043) | 22,686 (2,222) |
| Multiplicity | 15.0 (16.4) | 6.6 (6.0) | 6.9 (6.6) |
| Completeness (%) | 99.86 (100.00) | 98.47 (99.97) | 99.50 (100.00) |
| Mean I/sigma (I) | 19.7 (6.0) | 17.6 (4.3) | 26.2 (4.0) |
| R-merge | 0.114 (0.507) | 0.104 (0.377) | 0.017 (0.384) |
| R-pim | 0.0297 (0.1287) | 0.0434 (0.1664) | 0.0273 (0.1603) |
| $CC_{1/2}$ | 0.998 (0.942) | 0.997 (0.923) | 0.998 (0.875) |
| Wilson B factor ($Å^2$) | 21.98 | 22.10 | 24.99 |
| *Refinement* | | | |
| Reflections used in refienment | 15,395 (1519) | 30,227 (3,043) | 22,668 (2,222) |
| Reflections used for R-free | 715 (68) | 1,477 (143) | 1,152 (110) |
| R-work | 0.1915 (0.2088) | 0.2034 (0.2136) | 0.1975 (0.2722) |
| R-free | 0.2419 (0.2628) | 0.2556 (0.3023) | 0.2591 (0.3327) |
| Number of non-hydrogen atoms | 1837 | 4205 | 2159 |
| Macromolecules | 1726 | 3939 | 2013 |
| Solvent | 101 | 266 | 146 |
| Protein residues | 228 | 522 | 256 |
| RMS bonds (Å) | 0.007 | 0.008 | 0.007 |
| RMS angles (°) | 0.77 | 0.95 | 0.89 |
| Ramachandran | | | |
| Favoed/allowed/outliers (%) | 97.73/2.27/0.00 | 96.72/2.70/0.58 | 98.02/1.98/0.48 |
| Clashscore | 4.62 | 3.49 | 3.74 |
| Average B-Factor | 24.36 | 28.80 | 34.29 |

* Values in parentheses are for the highest resolution bins

**Table S6.** Idealized geometries and allowed deviations for catalytic constellation

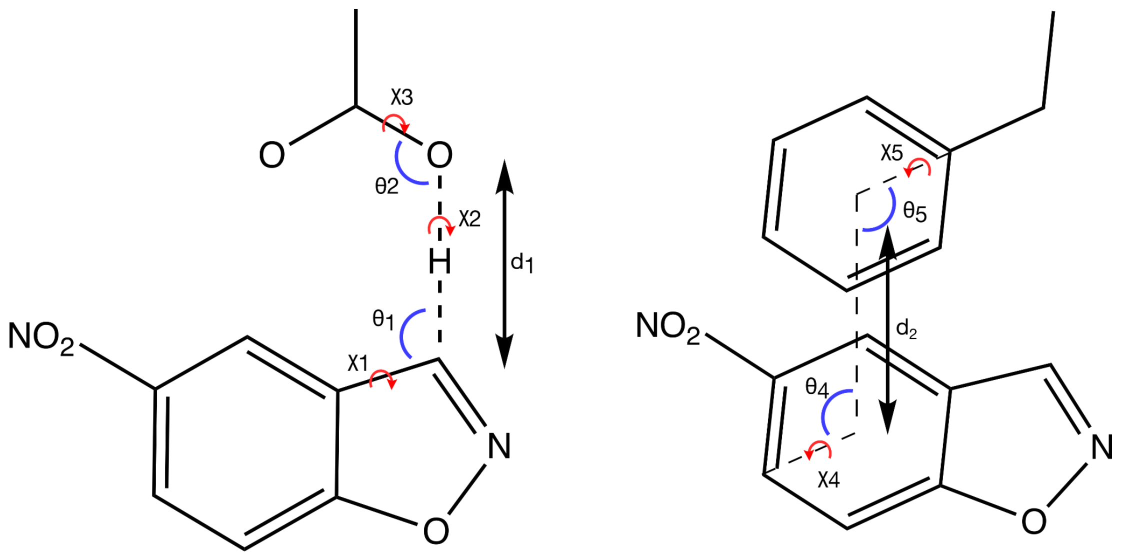

| | optimal distance/angle (Å/°) | deviation (Å/°) | Periodicity (°) |
|---|---|---|---|
| $d_1$ | 2.93 | 0.2 | (covalent) |
| $\theta_1$ | 127.6 | 5.0 | 360 |
| $\theta_2$ | 120.0 | 5.0 | 360 |
| $\chi_1$ | 180.0 | 7.0 | 360 |
| $\chi_2$ | 0.0 | 0.0 | 5 |
| $\chi_3$ | 180.0 | 10.0 | 360 |
| $d_2$ | 4.5 | 0.2 | |
| $\theta_3$ | 90 | 5.0 | 360 |
| $\theta_4$ | 90 | 5.0 | 360 |
| $\chi_4$ | 90 | 15.0 | 180 |
| $\chi_5$ | 90 | 15.0 | 180 |

**Table S7.** Protonation patterns of histidine residues for the systems simulated in this work.

| System | Non-Standard Protonation States |
|---|---|
| Des27 | HIE153, HIP164 |
| Des27.7 | HID92, HIE153, HIP164 |

**Table S8.** Non-standard force field parameters used to describe the substrate 5-nitrobenzisoxazole in our conventional molecular dynamics simulations.[a]

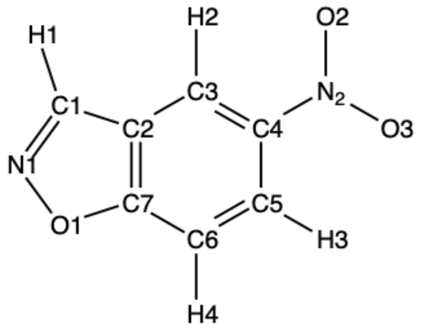

| Atom Name | Atom Type | Charge |
|---|---|---|
| O1 | os | -0.199614 |
| N1 | nd | -0.313448 |
| C1 | cc | 0.154120 |
| H1 | h4 | 0.151279 |
| C2 | ca | -0.050761 |
| C3 | ca | -0.250849 |
| H2 | ha | 0.234663 |
| C4 | ca | -0.010267 |
| N2 | no | 0.742907 |
| O2 | o | -0.450408 |
| O3 | o | -0.450408 |
| C5 | ca | -0.070430 |
| H3 | ha | 0.198146 |
| C6 | ca | -0.432848 |
| H4 | ha | 0.229481 |
| C7 | ca | 0.518437 |

[a] All parameters were obtained using the General AMBER Force Field 2 (GAFF2) (*62*), as outlined in the Experimental Section in the main text.

**Table S9.** List of neutralized residues and histidine protonation patterns used in our EVB simulations of Des27 and Des27.7 variants, in complex with substrates 5-nitrobenzisoxazole. [a]

| Residue Type | Residue number |
|---|---|
| Asp | 25, 64, 68, 98, 107, 246 |
| Glu | 13, 19, 26, 29, 66, 71, 77, 89, 97, 100, 206, 223, 248, 253 |
| Lys | 28, 74, 78, 101, 222, 226, 249 |
| Arg | 30, 33, 38, 57, 67, 229, 243, 252 |
| His-δ | 92 |
| His-ε | 153 |

[a] Shown here are the residues that fall outside the explicit simulation sphere and were thus kept in their neutral form to avoid system instabilities created by having charged residues outside the water droplet (this is standard practice for such simulations). All other residues were kept in their usual ionization state at physiological pH. In the case of the histidine side chains, His-ε and His-δ indicate histidine side chains protonated at the $N_{\varepsilon 2}$ and $N_{\delta 1}$ nitrogen atoms, respectively.
